# Hyperactivity preferentially disrupts scotopic retinal circuits in the early stages of diabetic retinopathy

**DOI:** 10.64898/2026.11.05.764529

**Authors:** Sheba Annie Philip, Narendra Pratap Singh, Priyanka Parida, Harikrishna Kakkatt, Aravind Swaminathaprasath, Lijitha Krishna Padikkappurath, Sejal Kumaravel, Santhosh Sethuramanujam

**Author notes:** Correspondence: Santhosh Sethuramanujam, Department of Biotechnology, Bhupat and Jyoti Mehta School of Biosciences, Indian Institute of Technology Madras, Chennai – 600036, India. These authors contributed equally (Narendra Pratap Singh, Priyanka Parida, Harikrishna Kakkatt).

## Abstract

Diabetic retinopathy is a leading cause of blindness worldwide. In this disease, visual impairments result from neurovascular dysfunction in the retina. However, the specific retinal circuits and the mechanisms underlying their dysfunction remain unclear. Here, in mouse models of diabetes, we evaluated scotopic (night vision) and photopic (day vision) retinal circuit function in the early stages of the disease. Using a combination of immunohistochemical, behavioral and electrophysiological techniques, we show that vision impairments early in the disease are not associated with widespread degeneration of the retinal circuitry but with abnormalities localized to scotopic circuits. Specifically, we observed swelling of axon terminals in rod but not cone bipolar cells. Importantly, these abnormalities were closely linked with bipolar cell activity, as rod bipolar cells showed higher intrinsic activity than cone bipolar cells, and inducing hyperactivity in control retinas was sufficient to recapitulate the pathology in diabetic rod bipolar cells. Further, the swollen rod bipolar cell terminals exhibited an increase in oxidative stress, consistent with cellular dysfunction. In line with this notion, electrophysiological measurements demonstrated that the contrast sensitivity of retinal ganglion cells was preferentially attenuated under scotopic conditions. Finally, behavioral experiments showed that visual sensitivity under scotopic conditions was affected earlier in disease progression compared to photopic conditions. Together, these findings reveal that scotopic circuits are preferentially affected in the early stages of diabetic retinopathy and that hyperactivity is sufficient to drive these abnormalities.

## Introduction

Diabetic retinopathy is a common complication of diabetes, which can cause progressive vision loss and, in severe cases, blindness. An estimated 100 million people worldwide are affected by this eye disease, with both India and the United States reporting around 10 million cases each^1–3^. Although traditionally considered a microvascular disease, mounting evidence in both humans and rodent models indicates that vision impairments can result from neurodegeneration in the retinal circuitry, which occurs in parallel with the vascular changes^4–10^. For instance, it has been reported that there is a marked reduction in retinal thickness and increased expression of apoptotic markers, indicating increased neuronal death in the disease^5,11–17^. Further, studies reveal attenuation of signals in electroretinogram measurements, indicating that circuit function in the retina is affected^12,14,17–21^. However, it remains unclear whether specific neuronal circuits are preferentially affected in the disease, and what mechanisms underlie their dysfunction.

Retinal circuits can be divided into photopic and scotopic circuits encoding day vision and night vision, respectively. Photopic circuits consist of cone photoreceptors and cone bipolar cells (CBCs), which in turn are connected to retinal ganglion cells, the output neurons of the retina (**Figure 1A**). In contrast, the primary scotopic circuits comprise rod photoreceptors and rod bipolar cells (RBCs), which do not directly connect to ganglion cells. Instead, they connect to the CBCs via the AII amacrine cell **(Figure 1A)**. The scotopic neurons have a variety of specializations that facilitate the reliable transmission of single-photon responses^22–24^, a critical feature of night vision. In contrast, photopic neurons are geared to respond faster with high spatial and chromatic acuity in day vision^25,26^. Given the distinct properties of these neurons, it is not surprising that visual disorders tend to preferentially affect neurons that encode either night vision^27,28^ or day vision^29,30^. Notably, dysfunction begins in specific neuronal subpopulations, but disease progression can lead to neuronal dysfunction in other cell types^27^. In the case of diabetic eye disease, most studies have tended to focus on the disease stages where widespread degeneration in the circuitry is prominent^5,11,12,14,15,31–33^. Alternately, studies have focused on the scotopic pathway, making it difficult to determine whether photopic circuitry is altered at these stages^34–38^. Given these limitations, it has remained unclear whether diabetes preferentially affects photopic or scotopic circuits.

**Figure 1:**
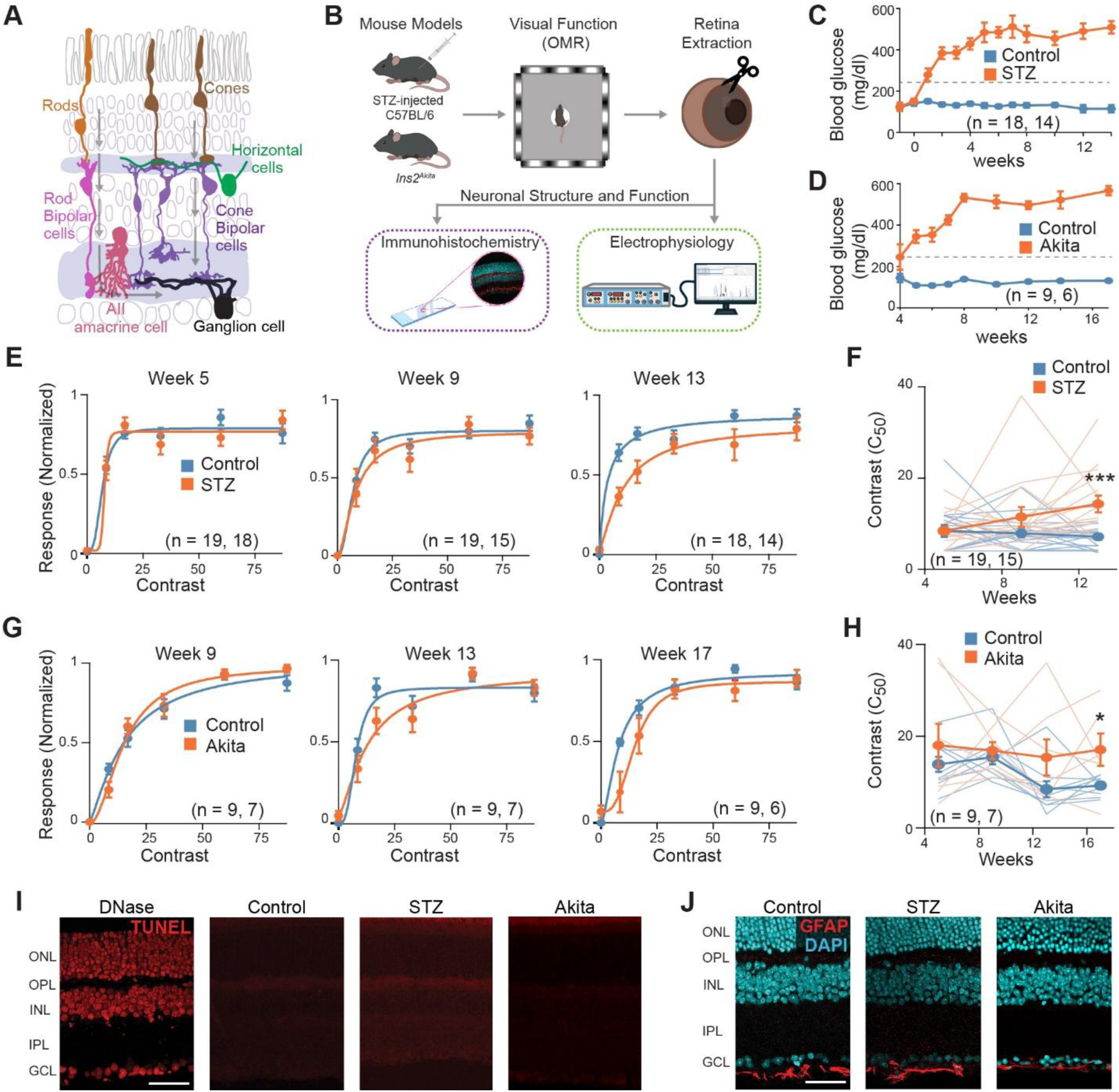
Vision impairments in diabetic mouse models preceded classical neurodegenerative markers. **(A)** Schematic of a retinal cross-section illustrating the organization of neuronal cell types within the scotopic and photopic pathways. Photoreceptors (rods and cones) transmit visual signals to bipolar cells. The arrows indicate the flow of information in the two pathways. Rod signals are relayed through rod bipolar cells (RBCs) to AII amacrine cells, which subsequently transmit the signal to cone bipolar cells (CBCs) and ultimately to retinal ganglion cells (RGCs). Cone signals are transmitted to RGCs directly through CBCs. **(B)** Schematic of the experimental workflow. STZ-injected and Ins2^Akita^ mice were assessed for visual impairment using the optomotor response (OMR) assay, followed by retina extraction for immunohistochemical and electrophysiological analyses. **(C)** Line graph showing blood glucose levels in control (blue) and STZ-injected (orange) mice over the experimental period. Following STZ administration, blood glucose levels in STZ-injected mice exceeded the hyperglycaemia threshold of 250 mg/dL (grey dashed line). Data represented as mean ± SEM. **(D)** Same as (C), showing blood glucose levels in control and Ins2^Akita^ mice. Note the change in the x-axis. **(E)** The contrast response function of OMR behavior (See Methods; **Figure S2**) under ambient light conditions in STZ-induced diabetic animals (orange) compared to control (blue). The responses were evaluated at 5, 9, and 13 weeks post-STZ injection. The data were fitted to the Naka-Rushton equation, as shown by the solid line. Data represented as mean ± SEM. **(F)** A comparison of the half-maximal contrast response (C_50_) values (See Methods) derived from the Naka-Rushton fits in STZ-induced diabetic animals compared to control at the timepoints shown in **E**. The population average is shown by the circles, and individual animal responses are shown by the thin lines. The STZ-diabetic animals had significantly higher C_50_ at 13 weeks post-STZ injection (***p < 0.001, n = 18 in the control group and 14 in the STZ group). Data represented as mean ± SEM. **(G, H)** Similar to E and F, but a comparison of the contrast sensitivity in the diabetic Akita models compared to littermate controls. Note that the time points differed from those in the STZ experiments. The Akita animals had significantly higher C_50_ at 17 weeks after birth (*p = 0.03, n = 9 in control and 6 in Akita group). **(I)** Representative TUNEL staining in mouse retinal cross sections in control, STZ and Akita animals, compared to a DNase-I-treated positive control. TUNEL-positive cells were minimal in both diabetic models. Scale bar: 40 µm **(J)** Representative retinal sections from a control, STZ-injected and Ins2^Akita^ mice showing GFAP expression (in red). GFAP expression was limited to the inner limiting membrane below the GCL, without extending into the outer retinal layers. The retinal layers are clearly demarcated by DAPI staining in blue. Scale bar: 40 µm.

What properties determine the propensity of a neuron to become dysfunctional in diabetes? As diabetes is a metabolic disorder, differences in the energy demand of neurons may influence vulnerability. For instance, higher energy demand may lead to cellular dysfunction due to oxidative stress, particularly in diabetic neurons^10,39–42^. A major determinant of energy requirements in neurons is synaptic activity and its associated homeostatic mechanisms^40,43,44^. Hence, strenuous neuronal activity could be a factor in determining a neuron’s vulnerability to dysfunction in diabetic retinopathy. Previous studies have shown that neurons in the scotopic circuitry exhibit hyperactivity in the early stages of diabetes^34,35,37^. However, it remains unknown whether RBCs and CBCs differ in intrinsic activity under physiological conditions and whether hyperactivity can drive dysfunction

Here, we examined the early stages of diabetic retinopathy in mouse models and found marked deficits in visual sensitivity in the absence of widespread degeneration in the retinal circuitry. These impairments were strongly associated with abnormalities in the scotopic but not photopic circuitry. For instance, the RBCs exhibited enlargement of their axon terminals, while the CBCs did not. Further, RBCs exhibited intrinsically higher activity than CBCs, and induction of hyperactivity in control retina recapitulated the enlarged RBC terminals, suggesting that high intrinsic activity may predispose RBCs to dysfunction. The enlarged RBC terminals in diabetic retina exhibited oxidative stress, consistent with cellular dysfunction. In line with RBC dysfunction, reduced visual sensitivity was observed under night-like conditions, apparent both at the level of retinal circuits and in the visual behavior of these mice.

## Results

### Vision impairments precede extensive retinal degeneration in diabetic mice

We began by evaluating the changes observed in the early stages of two well-established mouse models of diabetes. In the first model, diabetes was induced using streptozotocin (See Methods), which triggers apoptosis in pancreatic beta cells, leading to increased blood sugar levels within a week post-injection (**Figure 1C**). In the second model, diabetes manifests due to a genetic mutation that leads to misfolding of insulin, and we observed elevated blood sugar levels between 4 – 6 weeks after birth^45^ (**Figure 1D**). In both cases, the blood sugar levels were consistently high upto 13 – 17 weeks, which was the end point of the experiment (**Figure 1C, D**) (Control_STZ_: 133 ± 10 mg/dl, n = 18; Diabetic_STZ_: 455 ± 32 mg/dl, n = 14; Control_Akita_: 131 ± 4 mg/dl, n = 9; Diabetic_Akita_: 566 ± 22 mg/dl, n = 6; p < 0.001 for both models). These increased blood sugar levels were not associated with changes in body weight over the experimental period (**Figure S1**). To determine whether diabetic mice developed vision impairments, we monitored the contrast sensitivity in ambient light conditions **(Figure 1B)**. Here, we monitored the contrast sensitivity of the optomotor response, which is characterized by reflexive head movements in the direction of stimulus motion, a well-established marker in these mouse models^17,20,21,33^ (**Figure 1E - H**) (**Figure S2**; See Methods). Responses were quantified as the percentage of time the animal spent making head movements in the direction of stimulus motion (**Figure S2**). Gradually increasing the contrast of the visual scene results in a monotonic response that is well-fitted with a sigmoidal curve (**Figure 1E**). The contrast that elicited 50% of the peak response (C_50_) was considered an estimate of sensitivity (**Figure 1F**; See Methods). Monitoring the contrast sensitivity in control and diabetic animals revealed that the contrast sensitivity was significantly attenuated after 13 weeks post-induction in the STZ animals (control_STZ_ C_50_: 7.1 ± 0.7, n = 17; Diabetic_STZ_ C_50_: 14.3 ± 1.8, n = 16; p = 0.0012; **Figure 1E, F**) and 17 weeks after birth in Akita animals (Control_Akita_ C_50_: 9.2 ± 0.7, n = 9; Diabetic_Akita_ C_50_: 17.1 ± 3.5, n = 6; p = 0.029; **Figure 1G, H**). Given that it takes ∼4 weeks post-birth for hyperglycemia to develop in Akita mice, contrast sensitivity appears attenuated in both models after ∼3 months of hyperglycemia.

To pinpoint the retinal abnormalities that lead to vision impairments, we examined whether previously shown neurodegenerative markers manifest at this time point in diabetic retina (**Figure 1B**). Previous studies have shown increased apoptosis of third-order neurons in the diabetic retina^5,13,14^ (**Figure 1I**). Here, we monitored apoptosis levels using TUNEL staining in retinal cross-sections and found that the number of TUNEL-positive nuclei was not different between the diabetic models compared to their respective controls. Next, we examined an indirect marker of neurodegeneration, namely GFAP expression in Muller cells (**Figure 1J**). Under normal conditions, GFAP expression is largely confined to the end-feet processes of Müller cells in the inner limiting layer, but during diabetes induced gliosis, it extends further throughout the cell length, into the outer retinal layers^14,46^. We did not see an increase in GFAP fluorescence in the IPL in either model of diabetes compared to control, indicating an absence of gliosis in both models (**Figure 1J**). Finally, we evaluated the thickness of retinal layers, which has been shown to exhibit significant thinning, especially in the IPL^12,13^. Here, we focused on the central and mid-peripheral regions of the retina to ensure reliable measurements, as the natural tapering of the retina toward the periphery could confound our results (**Figure S3**). In line with the notion of a lack of widespread degeneration, there was no difference in the thickness of the synaptic or nuclear layers between the control and diabetic retina (**Figure S3**). Hence, these experiments clearly show that vision impairments in diabetic models occur before the onset of extensive neurodegeneration in the retinal circuitry.

### Rod bipolar cell terminals are preferentially altered in the early stages of diabetic retinopathy

The lack of widespread degeneration suggested that we were at an early stage of disease progression, where specific neuronal subpopulations could be preferentially affected. A preliminary screening using immunohistochemical labelling with well-established cell-specific markers revealed that RBCs showed the most prominent abnormalities in the diabetic retina (data not shown). RBCs project their axons into the innermost sublamina of the IPL, where they branch into multiple terminal endings, while their dendrites extend into the OPL to form synapses with rod photoreceptors (**Figure 2A**). The dendrites and somas of diabetic RBCs did not exhibit overt morphological differences compared to control (**Figure 2A – C**). In contrast, the axon terminals of diabetic RBCs in both models showed pronounced enlargement (**Figure 2A–C**). An analysis of the terminals revealed that the average size of RBC terminals was 31.5% larger in the STZ-injected retina and 38.7% larger in Akita retina compared to their corresponding controls (Control_STZ_: 7.8 ± 0.5 µm^2^, Diabetic_STZ_: 10.2 ± 0.5 µm^2^, n = 5 mice per group, p = 0.006; Control_Akita_: 7.5 ± 0.4 µm^2^, Diabetic_Akita_: 10.4 ± 0.8 µm^2^; n = 6 mice per group; p = 0.007; **Figure 2D**; See **Figure S4** and methods for details on analysis). This enlargement was seen in both the center and mid-periphery regions of the diabetic retina (**Figure S4**).

**Figure 2:**
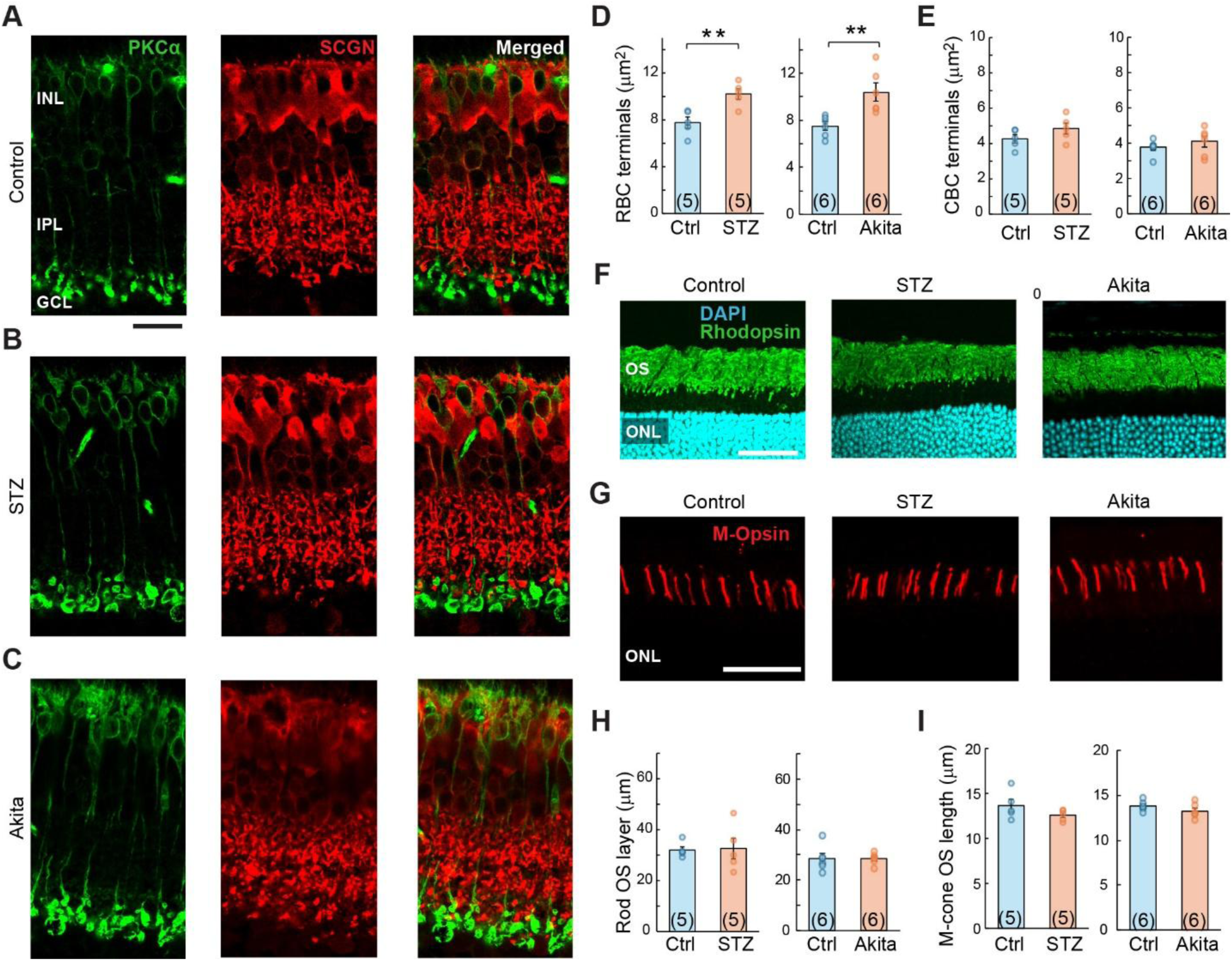
Rod bipolar cell terminals are preferentially affected in diabetes. **(A)** A representative retinal section from a control mouse labelled with RBCs (anti-PKCα in green) and a subset of CBCs (anti-secretagogin in red). Scale bar: 20 µm. **(B, C)** Similar to A, but for retinal sections in STZ induced and Akita diabetic mice. Notice that the RBC terminals are enlarged in both diabetic models. **(D)** Bar graph comparing the average size of RBC terminals between control and diabetic groups. Data represented as mean ± SEM, with individual data points shown as circles (n_STZ_= 5, n_Akita_= 6, **p < 0.01). **(E)** Bar graph comparing the average size of CBC terminals between control and diabetic groups. Data represented as mean ± SEM, with individual data points shown as circles (n_STZ_= 5, n_Akita_= 6). The analysis of CBC terminals was restricted to the layer adjacent to the RBC terminals, where individual terminals are easily demarcated (**Figure S4**). **(F)** Representative retinal sections from control, STZ-injected and Akita mice stained with DAPI (blue) and anti-rhodopsin antibody (green). Scale bar: 40 µm. **(G)** Representative retinal sections from control, STZ-injected and Akita mice showing M-opsin expression (red) in the outer segments of photoreceptors. Scale bar: 40 µm. **(H)** Bar graph comparing the height of the layer with rhodopsin-positive outer segments between control and diabetic groups. Data represented as mean ± SEM, with circles indicating the individual data points (n_STZ_= 5, n_Akita_= 6). **(I)** Bar graph comparing the lengths of M-opsin-positive segments between control and diabetic retinas. Data represented as mean ± SEM, with circles indicating the individual data points (n_STZ_= 5, n_Akita_= 6).

To evaluate whether the changes in the RBCs are mirrored in the CBCs, we stained the retina with secretagogin, which labels a subset of CBCs^47^ (Types 2, 3, 4, 5, 6, and possibly Type 8) (**Figure 2A – C**). Among these cells, the terminals of putative type 6 CBCs were examined, as individual terminals were easily identifiable (**Figure S4**). The results showed no significant difference in CBCs of both models of diabetic retina compared to their corresponding controls (Control_STZ_: 4.3 ± 0.2 µm^2^, Diabetic_STZ_: 4.9 ± 0.3 µm^2^, n = 5 mice per group, p = 0.179; Control_Akita_: 3.8 ± 0.2 µm^2^, Diabetic_Akita_: 4.1 ± 0.3 µm^2^; n = 6 mice per group; p = 0.396; **Figure 2E**). Tellingly, in the same retinal cross-sections, while RBC terminals exhibit enlargement, the CBCs do not (**Figure 2A – C**). This observation clearly indicated that the enlargement is cell-type-specific, as both type 6 bipolar cell terminals and RBCs stratify adjacent to each other in the IPL.

Next, we examined whether other neurons in the rod/cone circuits are differentially afflicted in diabetes. To assess rod photoreceptor dysfunction, we examined outer segment length, which has been shown to shorten in other neurodegenerative disorders^27^ (**Figure 2F, H**). We also examined the morphology of their terminals using PSD95, which shows the typical horseshoe labelling of rod spherules (**Figure S5**). At the time point where RBC terminals exhibited enlargement, there was no significant difference in the length and terminal morphology of the rod photoreceptors in control and diabetic retina (Control_STZ_: 32 ± 1.3 µm, Diabetic_STZ_: 32.6 ± 4.0 µm, n = 5 mice per group, p = 0.883; Control_Akita_: 28.5 ± 2.0 µm, Diabetic_Akita_: 28.5 ± 0.9 µm; n = 6 mice per group; p = 0.997; **Figure 2F, H; Figure S5**). This lack of change was mirrored in the cone photoreceptors, where the length of the outer segments—evaluated using M-opsin and PNA—was not significantly different between control and diabetic retina (Control_STZ_: 13.6 ± 0.7 µm, Diabetic_STZ_: 12.6 ± 0.3 µm, n = 5 mice per group, p = 0.195; Control_Akita_: 13.9 ± 0.2 µm, Diabetic_Akita_: 13.3 ± 0.3 µm, n = 6 mice per group, p = 0.185; **Figure 2G, I; Figure S5**). Further, the morphology of the cone pedicles outlined in the PNA staining were also not significantly different in diabetic retina compared to control (**Figure S5**). Hence, these experiments clearly reveal that the outer retina is relatively unaffected at these early stages, and the abnormalities are localized to the RBC terminals in the inner retina.

### Hyperactivity leads to enlargement of rod bipolar cell terminals

In retinal neurons, several morphological changes have been reported to correlate with neurodegeneration. For instance, studies show changes in dendritic architecture and withdrawal^27,48^. There is also evidence of dendritic swelling in the diabetic retina^49^. Our experiments indicate a new marker for the early stages of diabetic retinopathy progression, in the form of localized swelling in the axon terminals of RBCs. More importantly, these changes were not observed in corresponding CBC terminals. This preferential affliction led us to examine potential mechanisms underlying swelling of RBC, but not CBC terminals.

It has previously been suggested that the shape of RBC terminals is adaptable and linked to neuronal activity^50^. Hence, it is plausible that neuronal/synaptic activity is linked to terminal swelling. We began by testing whether the RBCs and CBCs have different levels of neuronal activity. Towards this endeavour, we monitored the expression of cFos, an immediate-early gene, which is transiently activated during increased neuronal activity^51,52^. Our experiments revealed that cFos expression was higher in RBCs compared to the adjacent CBCs in the control retina (RBC = 32.0 ± 4.7 a.u., CBC = 11.1 ± 2.6 a.u., n = 5 per group, p = 0.009; paired t-test; **Figure 3A, C;** See **Figure S6** for details on the separation of RBCs and CBCs). This difference was further exacerbated in the diabetic retina, where a large population of RBCs exhibited increased cFos expression (RBC = 68.1 ± 7.9 a.u., CBC = 30.9 ± 10.9 a.u., n = 5 per group, p = 0.015; paired t-test; **Figure 3B - D**). Hence, RBCs inherently exhibit higher neuronal activity in both physiological and pathological conditions.

**Figure 3:**
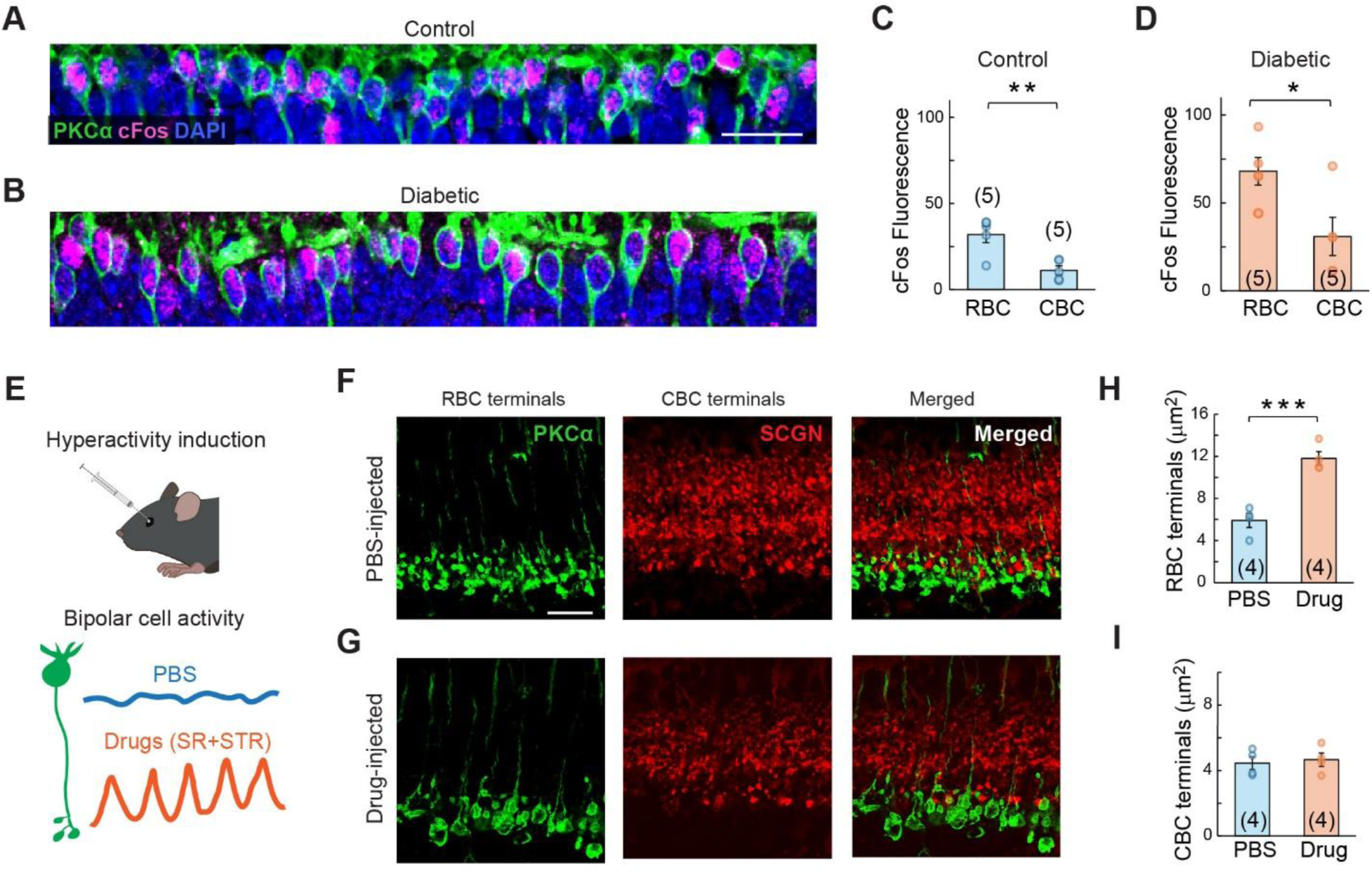
Hyperactivity enlarges rod bipolar cell terminals. (**A, B**) Representative cross-sections of the retinal inner nuclear layer (INL) labelled for cFos (magenta) expression in RBCs (green), in control (A) and STZ-induced diabetic (B) animals. DAPI (blue) is used to label all nuclei in the INL. Scale bar: 20 μm. **(C)** A comparison of the mean cFos fluorescence intensity in RBC and CBC nuclei in the control retina (n = 5). See Figure S6 for the separation of RBCs and CBCs. cFos fluorescence was significantly higher in the RBCs compared to CBCs (**p = 0.009; paired t-test). The individual samples are represented as open circles and the bar represents mean ± SEM. **(D)** Similar to C, but for diabetic retina. As seen in control, cFos fluorescence in diabetic tissue was significantly higher in the RBCs compared to CBCs (*p = 0.015; paired t-test). **(E)** Schematic of hyperactivity induction in retina by intravitreal injection of the GABA receptor antagonist, gabazine (1mM), and the glycine receptor antagonist, strychnine (100µM). Intravitreal PBS injection was used as a control. The bottom panels indicate the likely fluctuations in membrane potential observed in RBCs due to these injections. **(F)** Retinal cross-sections of the inner plexiform layer showing RBC terminals (PKCα - green) and CBC terminals (secretagogin – red) in PBS-injected control retina. Scale bar: 20 µm. **(G)** Similar to (B) but for drug-injected hyperactive retina. (**H, I**) A comparison of the RBC (H) and CBC terminal area (I) in hyperactive and control retina. Hyperactivity significantly increased RBC terminal area (***p <0.001, n = 4 in each group), while not altering the CBC terminals (p = 0.725, n = 4 in each group). Data represented as mean ± SEM. The individual samples are represented as open circles.

To directly test whether increasing neuronal activity can lead to terminal swelling, we induced hyperactivity in the control retina. Previous studies have shown that diabetic RBCs show increased activity due to attenuation of feedback inhibitory circuits^34–37^. In line with this notion, we find that synaptic proteins in inhibitory neurons in this layer are altered in diabetic retina (**Figure S7**). Hence, we evaluated whether inducing hyperactivity in the control retina—by abolishing inhibitory neurotransmission—leads to terminal enlargement (**Figure 3E**). We blocked retinal inhibitory circuits by intravitreal injection of 1 mM gabazine and 100 µM strychnine, antagonists of GABAA and glycine receptors, respectively (See Methods). A single injection of the drug mixture was delivered intravitreally, and the enlargement of RBC and CBC terminals was observed after 3 – 7 days post-injection. The analysis was restricted to the central region of the retina because the effects were more pronounced there. We observed that RBC terminals in the hyperactive retina were twice the size of terminals in PBS-injected control retina (PBS: 5.9 ± 0.7 µm^2^; Drug: 11.8 ± 0.6 µm^2^; *n* = 4 retina per group; *p* < 0.001; **Figure 3F - H**). In comparison, the CBC terminals were not significantly altered in the hyperactive retina (PBS: 4.5 ± 0.4 µm^2^; Drug: 4.7 ± 0.4 µm^2^; *n* = 4 retina per group; *p* = 0.725; **Figure 3F, G and I**). These experiments indicate that global alleviation of inhibition leads to a preferential enlargement of RBC terminals, suggesting their propensity to be affected by neuronal activity.

### Diabetic rod bipolar cell terminals exhibit increased oxidative stress

Having established that RBC terminals are specifically enlarged in the early stages of diabetes, we examined whether this enlargement affected their function. Towards this endeavour, we examined oxidative stress in these terminals, a hallmark of diabetic retinopathy that is well-correlated with attenuated function^41,42^. We examined the expression of MnSOD (manganese superoxide dismutase) in diabetic terminals (**Figure 4**). MnSOD is an antioxidant enzyme located in the mitochondrial matrix that neutralizes superoxide radicals generated during oxidative phosphorylation. Hence, altered MnSOD levels are considered a marker of mitochondrial oxidative stress^53^. Upon visual inspection of the MnSOD expression in RBC terminals, it became evident that many of the enlarged terminals in diabetic retina lack distinct MnSOD puncta, which typically appear as small, bright spots (**Figure 4A, C, E**). Analysing the MnSOD puncta after dividing the terminals based on size revealed that terminals over 30 µm^2^ had a significantly lower MnSOD fluorescence compared to the smaller size terminals (20 – 30 µm^2^) (small terminals: 1.11 ± 0.02; large terminals: 0.96 ± 0.03; p = 0.0021; 5 mice per group; **Figure 4D**). To enhance our accuracy of detecting MnSOD in RBC terminals, we confirmed these observations using super-resolution imaging. The super-resolution system was able to reliably identify the localization of the MnSOD puncta to within 70nm which is more than twice the resolution of the confocal microscope (**Figure S8**). Analysis of the data acquired from super-resolution indicated that large terminals in diabetic retina had ∼25% lower MnSOD fluorescence compared to smaller size terminals (small terminals: 1.01 ± 0.08; large terminals: 0.74 ± 0.07; *p =* 0.043; 5 mice per group; **Figure 4E, F**). A number of controls indicated that our observations are likely due to localized changes in the diabetic retina. First, the expression of MnSOD in the population of RBC terminals was comparable in control and diabetic retina (**Figure 4B**). Second, the relative expression of MnSOD in the RBC layer compared to the rest of the IPL did not change in diabetic retina (**Figure S8**). Finally, we observed large puncta in the OPL (presumably mitochondria in the rod photoreceptor terminals), which remained unchanged in size (**Figure S8**). Hence, these experiments indicate that oxidative stress is specifically increased in enlarged RBC terminals in the early stages of disease progression.

**Figure 4:**
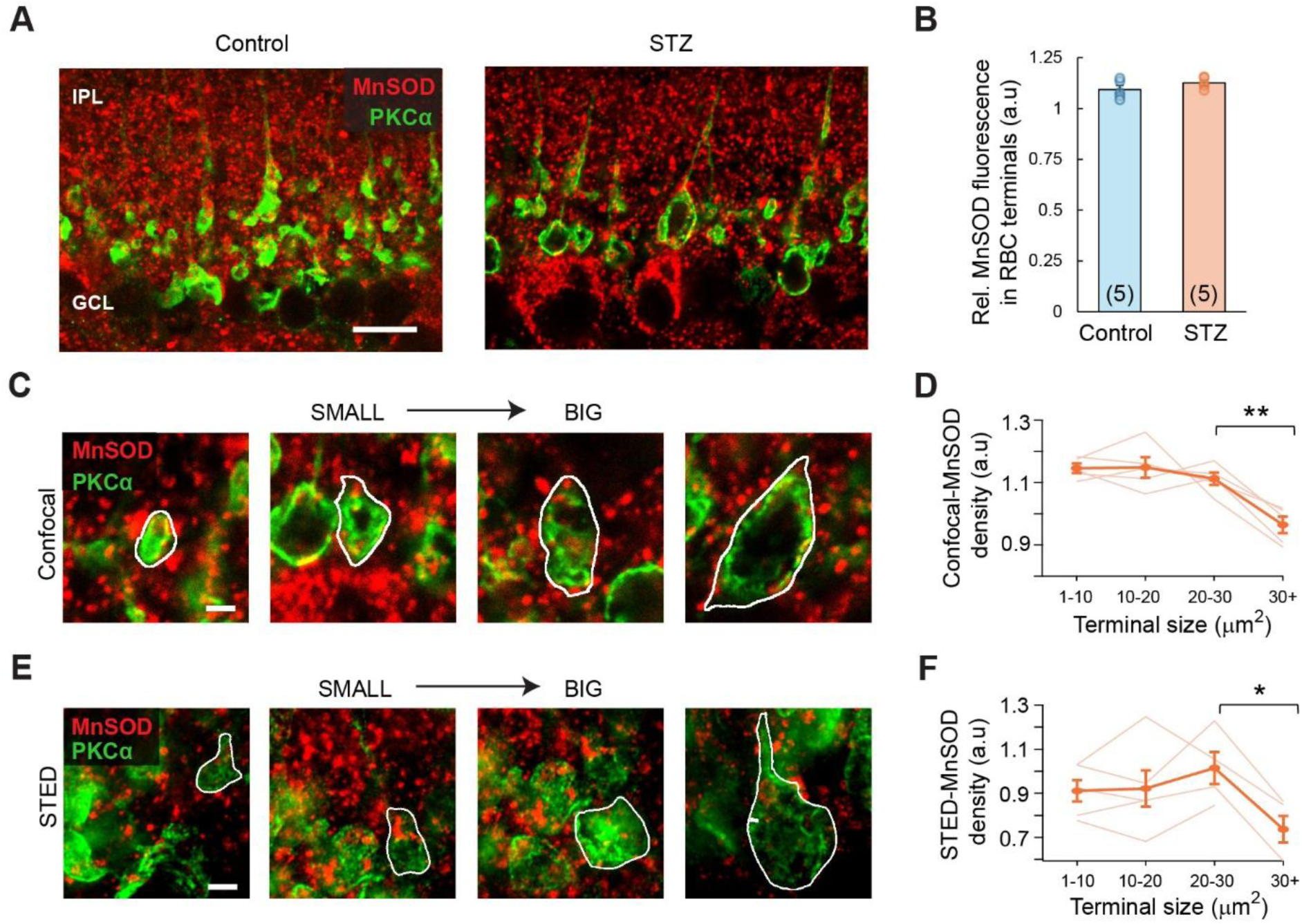
Oxidative stress manifests in the enlarged RBC axon terminals in the diabetic retina. **(A)** Retinal sections from control (left) and diabetic (STZ; right) mice labelled with anti-PKCα (in green) and anti-MnSOD antibody (in red). Scale bar: 10 µm. **(B)** Bar graph comparing the relative intensity of MnSOD fluorescence inside the terminals between control and diabetic retinas. See methods for details of the analysis. Data represented as mean ± SEM, with circles indicating the individual data points (n = 5 mice per group). **(C)** Representative images of RBC axon terminals of increasing size (outlined in white) and immunolabeled with anti-PKCα (green) and anti-MnSOD (red) in diabetic retina, acquired using confocal microscopy. See methods for details on identifying individual terminals Scale bar: 2 µm. **(D)** Line graph showing changes in MnSOD expression across increasing terminal sizes for each diabetic animal (thin lines). The group mean is represented by the solid line, with error bars indicating the SEM. The MnSOD fluorescence was reduced for the largest RBC terminals (**p = 0.002; n = 5 animals). Note that the Y-axis does not start from zero to better visualize subtle changes. (**E, F**) Similar to C and D, but for images collected using STED microscopy. As seen in the confocal images, MnSOD fluorescence was reduced for the largest RBC terminals (*p = 0.043; n = 5 animals).

### Scotopic contrast sensitivity is preferentially affected in diabetic retinal ganglion cells

To determine whether the observed cellular changes affect RBC function, we monitored the spike activity of retinal ganglion cells in ex vivo whole-mount preparations using single-cell loose-patch recordings (**Figure 5A, 1B**). We hypothesized that RBC dysfunction in the early stages of diabetes should preferentially affect retinal ganglion cell function under scotopic conditions compared to photopic conditions. Here, ganglion cell function was evaluated by measuring their contrast sensitivity to static light spots centered on the soma (**Figure 5D**; See Methods). Ambient light levels were modulated with neutral-density filters, and we used high-mesopic (∼100 R*/rod/s) and low-mesopic (∼10 R*/rod/s) conditions to assess contrast sensitivity^54^. We used light levels that were only 1 log unit apart, as it was difficult to elicit robust responses at low mesopic light after exposing the tissue to traditional photopic light (∼1000 R*/rod/s). For simplicity, high-mesopic light is referred to as photopic, and low-mesopic as mesopic. As shown in previous studies, we observed strong surround inhibition in photopic conditions and significantly weaker surround inhibition in mesopic conditions^55,56^ (**Figure S9**). More importantly, the mesopic conditions used here are rod-dominated and become cone-dominated at the employed photopic light levels^54,55^.

**Figure 5:**
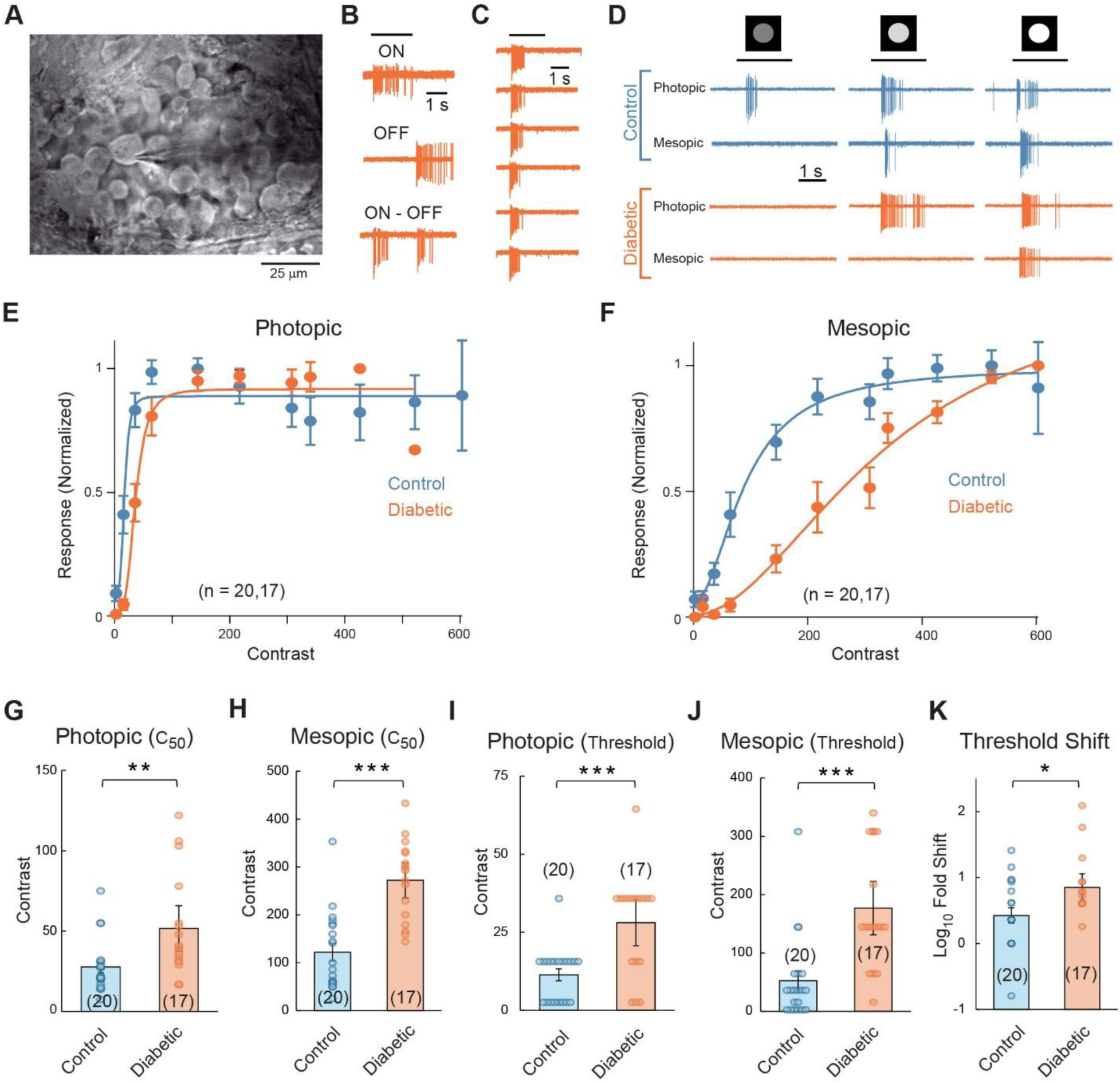
Diabetic insult preferentially impairs retinal ganglion cell contrast sensitivity under mesopic conditions. **(A)** Representative image showing loose patch configuration of a retinal ganglion cell in the wholemount preparation. **(B)** Light-evoked spike activity recorded from an ON, OFF and ON - OFF retinal ganglion cell in the diabetic retina. The responses to a static spot of light are shown. The duration of the light stimulus is indicated by the black line at the top. **(C)** Light-evoked spike responses of an ON ganglion cell in diabetic retina to a static spot of light over 6 trials. **(D)** Light-evoked spike activity recorded from an ON ganglion cell in a control retina compared to an ON cell in a diabetic retina, under photopic and mesopic conditions. The responses to a static spot of light at three different stimulus contrast levels are shown. The duration of the light stimulus is indicated by the black line at the top. **(E)** A comparison of contrast response functions of the light evoked spike activity of retinal ganglion cells in control (blue) and diabetic (orange) retinas, under photopic conditions. The data were fitted to the Naka-Rushton equation, shown by the solid line. Data represented as mean ± SEM. **(F)** Similar to E, but for mesopic conditions in the same population of retinal ganglion cells. **(G)** A comparison of the C_50_ values (from the Naka-Rushton equation fits) between control and diabetic retinas under photopic conditions. The individual cells are indicated by the circles, and the bar indicates the population average. The C_50_ was significantly higher in retinal ganglion cells in the diabetic retina compared to the control retina. (**p = 0.01; n = 20 in control and 17 in diabetic retina). Data represented as mean ± SEM. **(H)** Similar to G, but for the responses generated by mesopic conditions. As observed under photopic conditions, the C_50_ was significantly higher in diabetic ganglion cells compared to controls. (***p < 0.001; n = 20 in control and 17 in diabetic retina). Data represented as mean ± SEM. **(I, J)** A comparison of response threshold (see methods) in control and diabetic retinal ganglion cells under photopic (I) and mesopic (J) conditions. Threshold values were significantly higher in diabetic retinas under both light conditions. (Photopic: ***p = 0.001; mesopic: ***p < 0.001; n = 20 in control and 17 in diabetic retina). Data represented as mean ± SEM. **(K)** Comparison of the shift in response threshold (see methods) between control (n = 20) and diabetic (n = 17) retinal ganglion cells. Diabetic retinal ganglion cells exhibited a significantly greater threshold shift, indicating reduced contrast sensitivity compared to control retina (*p < 0.013; n = 20 in control and 17 in diabetic retina). Data represented as mean ± SEM.

In these experiments, we consistently recorded ON, OFF, and ON-OFF cells in both control and diabetic retina (**Figure 5B**). We did not attempt to further classify the subtypes, and responses from all cells were pooled for contrast sensitivity measurements. The data presented here are from 20 cells from control retina and 17 cells from the STZ-induced diabetic retina (4 animals each). On average, strong light stimulation elicited fewer spikes in diabetic retinal ganglion cells compared to their control counterparts (Control: 22 ± 3 spikes; diabetic: 13 ± 1 spikes; p = 0.003). Nevertheless, several physiological observations indicated a lack of widespread neurodegeneration in the diabetic retina. First, we observed robust spike activity across multiple repetitions of stimulation (**Figure 5C**). Second, we observed that complex computations were intact in the diabetic retina, as we observed both size selectivity and direction selectivity (**Figure S9**). Finally, we observed that the spontaneous activity of ganglion cells—known to increase in neurodegenerative disorders^57–59^—was less than 1 Hz in both control and diabetic retina (Control: 0.8 ± 0.1 spikes; Diabetic_STZ_: 0.1 ± 0.1 spikes).

Next, we evaluated contrast sensitivity using two parameters. First, we estimated the C_50_, the contrast that elicits a half-maximal amplitude, as shown in the visual behavior experiments in **Figure 1**. Second, we estimated visual threshold as the minimum contrast required to elicit spiking in the ganglion cell (See Methods). In photopic conditions, the C_50_ of control retinal ganglion cells was 27.7 ± 3.7, while that of diabetic ganglion cells was 51.7 ± 7.7 (**Figure 5D, E, G**). In the same set of ganglion cells, under mesopic conditions, the control ganglion cells had a C_50_ of 121.8 ± 18.2 while the diabetic ganglion cells exhibited a C_50_ of 272.1 ± 20.0 (**Figure 5D, F, H**). In both photopic and mesopic conditions, the observed C_50_ was significantly higher in diabetic ganglion cells compared to control ganglion cells (photopic C_50_: p = 0.01; mesopic C_50_: p < 0.001; **Figure G, H**). Similar effects were observed for visual threshold, where diabetic ganglion cells had a significantly higher threshold in both photopic and mesopic conditions (Control photopic: 11.4 ± 1.9, diabetic photopic: 28.1 ± 4.0, p = 0.001; Control mesopic: 52.6 ± 16.4, diabetic mesopic: 177 ± 25, p < 0.001; **Figure 5I, J**). Hence, diabetic ganglion cells had reduced contrast sensitivity in both photopic and mesopic conditions. However, the effect of diabetes on contrast sensitivity was more pronounced under mesopic conditions. This is best illustrated by comparing the shift in photopic sensitivity relative to mesopic sensitivity in individual ganglion cells. We observed that, in control ganglion cells, the mesopic threshold was 5 ± 1.3 fold higher than the photopic threshold, whereas in diabetic ganglion cells, the mesopic threshold was 15.9 ± 7.4 fold higher than the photopic threshold (p < 0.013; **Figure 5K**). Hence, the sensitivity of diabetic retinal ganglion cells was more adversely affected under mesopic than photopic conditions.

### Impairments in scotopic vision manifest earlier in diabetic mice

The experiments so far indicate that the anatomy and physiology of the retinal circuits underlying night vision are preferentially affected in the early stages of diabetic retinopathy. Finally, we evaluated whether these retinal changes are reflected in the vision impairments observed in diabetic animals. We predicted that the onset of vision impairments under scotopic conditions is likely to manifest earlier than under photopic conditions. To test this, we monitored the contrast sensitivity of the optomotor response in control and diabetic mice under both photopic and scotopic conditions (See Methods; **Figure 6**). The ambient light was modulated using neutral-density filters to achieve photopic (10^5^ R*/rod/s) and scotopic (1 R*/rod/s) conditions. As shown in previous studies, we observed a shift in the optomotor response profile under scotopic conditions compared with photopic conditions^60^ (**Figure S10**).

**Figure 6:**
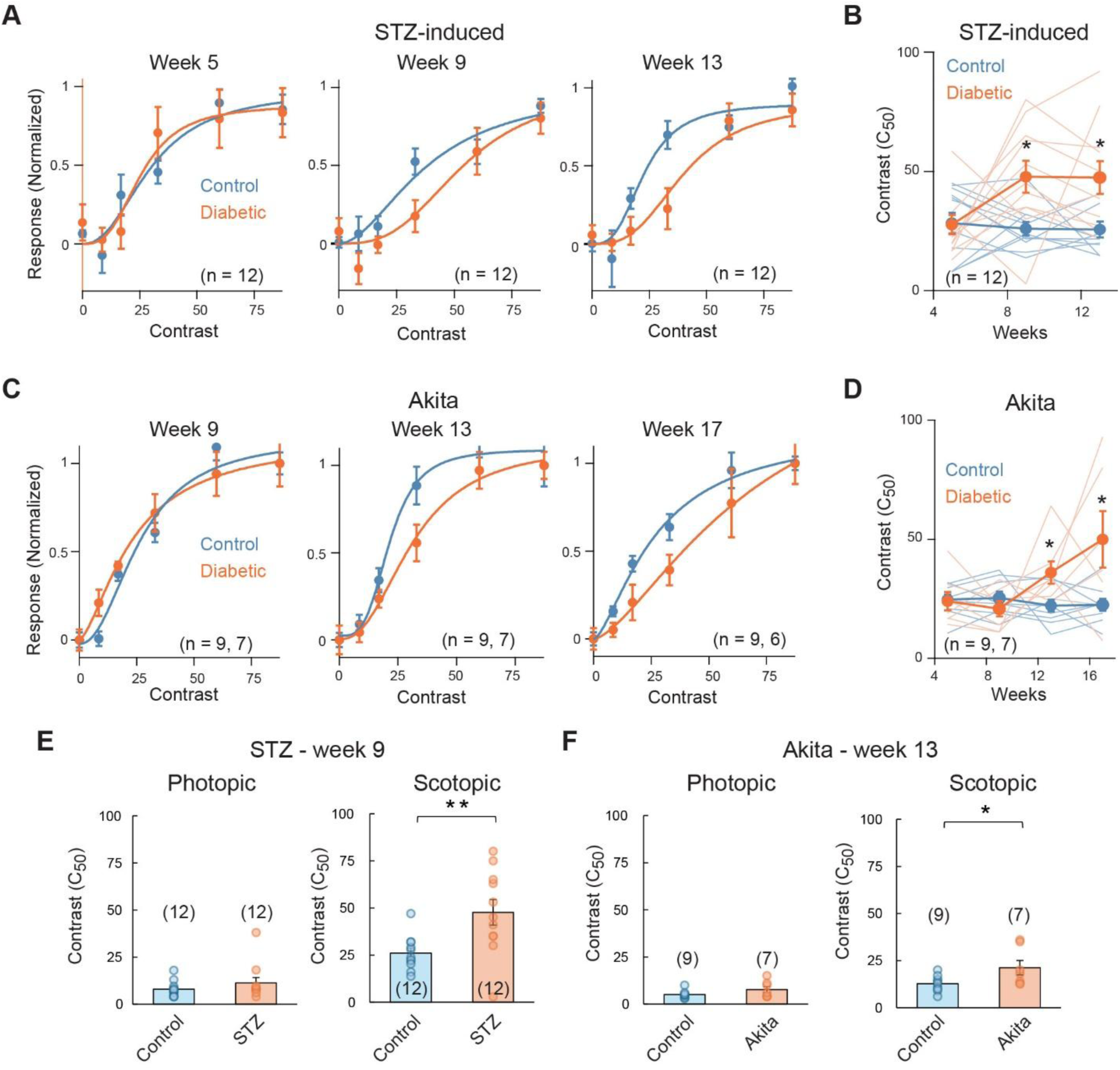
Contrast sensitivity deficits in diabetic mice under scotopic conditions. **(A)** The contrast response function of OMR behavior under scotopic conditions (See Methods) in STZ-induced diabetic animals (orange) compared to control (blue). The responses were evaluated at 5, 9, and 13 weeks post-STZ injection. The data were fitted with the Naka-Rushton equation depicted by the solid line. Data represented as mean ± SEM. **(B)** A comparison of the C_50_ values (See Methods) derived from the Naka-Rushton fits in STZ-induced diabetic animals compared to control at the timepoints shown in **A**. The population average is shown by the circles, and individual animal responses are shown by the thin lines. The STZ diabetic animals had significantly higher C_50_ at 9 weeks and 13 weeks post-STZ injection (*p = 0.01, n = 12 in each group). Data represented as mean ± SEM **(C)** Similar to A, but for Akita diabetic mice (orange) compared to wild-type littermates (blue) at 9, 13, and 17 weeks of age. **(D)** Similar to B, but for Akita diabetic mice compared to wild type littermates at 5, 9, 13, and 17 weeks of age. The Akita animals had significantly higher C_50_ at 13 and 17 weeks after birth (*p = 0.02, n = 9 in the control group and 7 in the Akita group). Data represented as mean ± SEM **(E)** A comparison of the C_50_ in control and STZ-induced animals in photopic (left) and scotopic (right) conditions, 9 weeks post-diabetic induction. The C_50_ of the STZ animals was significantly higher than that of the control only in scotopic conditions (**p = 0.008, n = 12 in both control and diabetic groups). Data represented as mean ± SEM **(F)** A comparison of the C_50_ in control and Akita animals in photopic (left) and scotopic (right) conditions, 13 weeks after birth. The C_50_ of the Akita animals were significantly higher than control only in scotopic conditions (*p = 0.021, n = 9 in the control group and 7 in the Akita group). Data represented as mean ± SEM

We monitored the scotopic contrast sensitivity of control and both models of diabetic mice at the same timepoints as observed for photopic conditions in Figure 1 (**Figure 6A, C**). Under scotopic conditions, the contrast sensitivity was significantly lower compared to photopic conditions (Photopic C_50_: 8.9 ± 1.7; Scotopic C_50_: 28.3 ± 4.3; n = 12; p = 0.002; **Figure 6E**). A comparison of control and diabetic animal behavior revealed that scotopic contrast sensitivity was significantly altered at 9 weeks post-induction in the STZ animals (control C_50_: 26 ± 2.8, n = 12; STZ C_50_: 47.7 ± 6.8, n = 12; p = 0.011; **Figure 6A, B**). The Akita animals show significantly reduced scotopic sensitivity 13 weeks after birth (control C_50_: 22.1 ± 2.5, n = 9; Akita C_50_: 36 ± 4.7, n = 7; p = 0.022; **Figure 6C, D**). In the same set of animals, at this time point, photopic contrast sensitivity was not significantly altered in both STZ (control C_50_: 8 ± 1.2, n = 12; STZ C_50_: 11.3 ± 2.9, n = 12; p = 0.307; **Figure 6E**) and Akita diabetic models (control C_50_: 8.4 ± 1.8, n = 9; Akita C_50_: 15.4 ± 3.9, n = 7; p = 0.126; **Figure 6F**). The photopic contrast sensitivity was significantly altered only after 13 weeks in the STZ model and 17 weeks in the Akita model (**Figure 1**). Hence, these results show that scotopic contrast sensitivity is affected 4 weeks earlier in the progression of diabetes than photopic contrast sensitivity.

## Discussion

The prevailing notion in diabetic retinopathy models is that vision impairments are engendered by widespread neurodegeneration in the retina. This is well-illustrated by numerous studies over the past 30 years, which show that visual impairments are strongly associated with widespread changes in the neuronal circuitry, even during the early stages of the disease^5,12–15,21,31,41,42,49,61–63^. Our experiments support an alternative model in which vision impairments in the early stages of the disease occur despite an apparent lack of indicators of widespread degeneration in the circuitry^36^. Multiple lines of evidence support this model. First, the traditional markers of neurodegeneration that indicate neuronal loss, including tissue thinning, apoptotic markers and gliosis, were largely absent. Second, morphological abnormalities were limited to specific subsets of neurons, consistent with localized dysfunction. Finally, functional evaluation of the circuitry using patch-clamp recordings clearly shows that a large fraction of the inhibitory circuits in the retina is predominantly intact. This is best evidenced by the observation of complex computations in the diabetic retina, including size-selectivity and direction-selectivity, which require precise excitatory and inhibitory inputs to ganglion cells. Hence, our findings clearly support the model that vision impairments—in the early stages of the disease—are not due to widespread changes but rather due to abnormalities confined to specific retinal circuits.

### Scotopic circuits are preferentially affected in the early stages of diabetic retinopathy

The central finding of this study is that scotopic vision and its underlying circuits are preferentially afflicted in the early stages of the disease. Previous studies evaluating photopic and scotopic vision have yielded mixed results; while most studies indicate that photopic vision is affected^14,17,20,21,33,42,62^, a few suggest that photopic vision is not affected but scotopic vision is^12,18^. One reason for such variability may arise from technical limitations such as use of anaesthesia in electroretinograms^64,65^. Alternately, variability could be due to differences in animal models or in the methodology for diabetic induction, as visual deficits begin at a wide range of time points, ranging from as early as 4 weeks^17,20^ post-diabetes to as late as 15 weeks^33^. Here, we overcome this variability by comparing photopic and scotopic contrast sensitivity deficits in the same set of animals and clearly show that scotopic contrast sensitivity in freely behaving mice was affected almost a month earlier than photopic contrast sensitivity in two diabetes models.

Importantly, our results provide insights into the circuit-level changes in the retina that could lead to abnormalities in scotopic vision by monitoring the contrast sensitivity of retinal ganglion cells. Previous studies have monitored the membrane and synaptic properties of RBCs and retinal ganglion cells. For instance, studies clearly reveal that diabetic insult increases spontaneous activity in RBCs^34–36^, likely due to reduced feedback inhibition^35,36^. Further, it has been suggested that the intrinsic and synaptic excitability of retinal ganglion cells could be increased in diabetes^37,57,66^. However, it was unclear how these abnormalities affect retinal ganglion cell function and whether they can explain the abnormalities observed in visual behaviour. Our results provide the missing link, showing that diabetic retinal ganglion cells exhibit reduced contrast sensitivity compared with ganglion cells under control conditions. Furthermore, these deficits are greater at mesopic than at photopic light levels. Given that low mesopic light levels are dominated by the scotopic pathway^54^, these observations indicate that neurons in the scotopic circuit are more affected by the disease.

Further evidence for preferential affliction of scotopic neurons is provided by morphological abnormalities in RBCs but not CBCs. Specifically, we observed a robust enlargement of RBC terminals in the diabetic retina. Such swellings are thought to arise from cytoskeletal abnormalities, which can affect intracellular transport^67^ and, consequently, attenuate neuronal function. Hence, there is a strong correlation between morphological abnormalities and visual function^27,48,68^. While previous studies in the diabetic retina have reported changes in the dendritic architecture and swelling of both axons and dendrites in ganglion cells^49,57^, our experiments reveal that swelling of RBC terminals is a robust early marker in the diabetic retina. Moreover, we observed a selective increase in the oxidative stress of these terminals, indicating dysfunction. Hence, the results presented here comprehensively show that scotopic neurons—particularly RBCs—and their function are affected in the early stages of diabetic retinopathy.

### Hyperactivity as a driver of dysfunction in diabetic rod bipolar cells

The experiments presented here provide evidence for the role of neuronal activity in contributing to dysfunction in RBC terminals. This is best illustrated by the proof-of-principle experiment, where inducing hyperactivity in bipolar cells in the control retina leads to swelling of their terminals, mimicking the effects observed in diseased terminals in the diabetic retina. Interestingly, even when inhibitory circuits were blocked in both RBCs and CBCs, the terminal enlargement was prominent only in RBCs. Hence, the RBC terminals may be inherently more susceptible to damage from hyperactivity. This is further compounded by the observation that RBCs exhibit higher neuronal activity than CBCs under both physiological and disease conditions. Hence, the tendency towards higher activity in RBCs could be a crucial determinant of the manifestation of diabetes-induced dysfunction in these neurons.

Several studies suggest a proclivity in RBCs towards hyperactivity. RBCs have low-voltage calcium channels that do not inactivate, unlike CBCs^69^. This indicates that RBC synapses will be activated at relatively hyperpolarized membrane potential for longer periods of time. Indeed, direct monitoring of glutamate release indicates that RBCs exhibit a more sustained release than CBCs^70^. Further, the RBCs are electrically coupled to the AII amacrine cells, which is known to cause oscillatory hyperactivity upon slight perturbation^71^. They also receive strong, sustained inhibition from feedback networks^72,73^ and attenuation of these circuits, as observed in diabetes, could further intensify activity^34–36^. Hence, the RBC is highly susceptible to increased activity due to both intrinsic and network properties. Surprisingly, the CBCs—despite being connected to the RBCs (via the AII amacrine cells)—do not seem to exhibit high activity. This is likely due to the observation that signal transmission from the AII amacrine cells to CBCs is weak^74,75^. Consequently, hyperactivity in the RBCs may not be transferred to the CBCs. This would explain the lack of increase in spontaneous activity at the level of retinal ganglion cells (but see^57,66^). Hence, the bipolar network properties may facilitate a preferential increase in activity in RBCs but not CBCs. Importantly, the increased activity is associated with dysfunction, likely due to oxidative stress in the diabetic terminals. Hence, it will be interesting to test whether controlling hyperactivity in diabetic retina prevents the manifestation of cellular dysfunction in RBCs.

The observations of this study may have implications beyond diabetic retinopathy, as several neurodegenerative disorders exhibit hyperactivity^76–78^. It will be interesting to observe whether hyperactivity develops in specific neuronal populations in these diseases and whether it is sufficient to induce dysfunction.

### Implications for diabetic retinopathy in humans

Whether observations made in rodent models of diabetes have implications for humans has long been debated. For example, neovascularization which is the hallmark of diabetic retinopathy in humans, is not readily observed in rodents, possibly because the short lifespan of rodents limits the extent of disease progression^79^. Nevertheless, as indicated earlier, several well-characterized features of the neural component of the disease observed in human patients are well-captured in rodent models^5,11,12^. Furthermore, RBCs and the neurons in the scotopic pathway, which are critical for encoding scotopic vision, are evolutionarily conserved in humans^80^. Hence, it is possible that scotopic vision is preferentially affected in diabetes, making it a potential diagnostic marker for identifying the early stage of the disease. While studies have focused on contrast sensitivity, visual acuity and colour deficits in human patients^7,8,81–85^, deficits in night vision remain an open question.

## Materials and Methods

### Animal maintenance

Male C57BL/6J and C57BL/6-*Ins2^Akita^*/J (Akita) mice (RRID:IMSR_JAX:003548) of both sexes (3-4 weeks old, weighing 15-20 g) were housed in the Institute Animal House at the Indian Institute of Technology Madras, India. All experimental procedures involving animals were approved by the Institutional Animal Ethics Committee of the Indian Institute of Technology Madras (# IITM/IAEC-1/SS-1/2024). The mice were maintained in cages under controlled temperature conditions (20 – 23° C) with a 12-hour light-dark cycle and had free access to food and water.

### Diabetes induction

Diabetes was induced in 2-month old C57BL/6J male mice by a series of low-dose streptozotocin injections (50 mg/kg daily for 5 consecutive days) delivered intraperitoneally. Prior to injection, the animals underwent a six-hour fast and were administered the drug in 0.1 M citrate buffer (40 mM sodium citrate, 60 mM citric acid, pH 4) within 10 minutes of dissolution. Age-matched control mice received injections of the vehicle buffer alone. Akita animals spontaneously developed diabetes between 4 – 6 weeks after birth. Blood glucose levels were measured using a glucometer (Accu-Chek) after a six-hour fast, by drawing blood from the animal’s tail tip. Mice having fasting blood glucose levels above 250 mg/dl were classified as diabetic.

### Optomotor response (OMR)

The OMR setup consisted of a central elevated pedestal surrounded by four identical monitors that displayed vertical black-and-white sine-wave gratings. The background illumination was set at 10^5^ photoisomerisations/s (R*/s) for photopic conditions and 1 R*/s for scotopic conditions. During each trial, the animal was placed on the pedestal and allowed to move freely while exposed to visual gratings rotating at a constant speed of 27°/sec. Head movements were recorded by a camera mounted on the top. Each trial presented gratings of a specific Michelson contrast for 60 seconds in the clockwise direction, followed by another 60 seconds in the counterclockwise direction. Michelson contrast is calculated as (I_max_-I_min_)/(I_max_+I_min_), where I_max_ is the maximum intensity of the grating, and I_min_ is the minimum intensity of the grating. The videos were analyzed using DeepLabCut^86^ to track the coordinates of the animal’s snout. Next, using these coordinates, the angular displacement of the snout relative to the center of the body was estimated with custom-written code in Python. Finally, we estimated the OMR response as the percentage of time the snout displacement is in the same direction as the stimulus. An OMR response of 50% indicates that the animal could not detect the stimulus, as it was moving its head randomly in either the clockwise or the anticlockwise direction, while any value significantly higher than 50% indicates that the animal could detect the stimulus.

### Single cell Patch Recordings

Animals were dark-adapted for 45 min, after which they were euthanized. The eyes were rapidly enucleated, and the retinas were carefully dissected in oxygenated Ringer’s solution under a stereomicroscope (Olympus SZ51) with red-light illumination. The isolated retina was flatmounted ganglion cell side up on a 0.22 µm membrane filter (Millipore) containing a pre-cut window to allow light to reach the photoreceptor layer. The mounted retina was transferred to a recording chamber and continuously superfused at 4ml/min with oxygenated Ringer’s solution containing (in mM): 110 NaCl, 2.5 KCl, 1 CaCl₂, 1.6 MgCl₂, 10 dextrose, and 22 NaHCO₃. The solution was continuously bubbled with 95% O₂ and 5% CO₂, and the temperature was maintained at 32–34°C.

RGCs were visualized using a real-time monochrome infrared video camera (Dage-MTI IR-2000) mounted on an upright fluorescent microscope (Olympus BX51WI) equipped with a 40× water-immersion objective. Loose patch spike recordings were obtained with 5–8 MΩ borosilicate glass pipettes filled with Ringer’s solution. Recordings were made with a Multiclamp 700B amplifier (Molecular Devices), digitized at 10 kHz with a Digidata 1550B (Molecular Devices) and viewed using pClamp 11. Spikes were detected and analyzed as described before using custom-written routines in MATLAB^87^.

Visual stimuli were generated using a digital light projector (Wzatco Yuva Plus; refresh rate: 60 Hz) and focused onto the photoreceptor outer segments through the microscope sub-stage condenser. The background illumination was set at 100 photoisomerisations/s (R*/s) for photopic conditions and 10 R*/s for mesopic conditions. Background intensities were adjusted using neutral density filters mounted on motorised filter wheels (Thorlabs). Stimuli were designed using StimGen, a custom presentation GUI written in Python with PsychoPy^88^ (https://github.com/benmurphybaum/StimGen). For contrast-response measurements, a 200 micron diameter static spot was used with varying Weber contrasts. Weber contrast is calculated as (I_stim_-I_bg_)/I_bg_, where I_stim_ is the intensity of the stimulus and I_bg_ is the intensity of the background. For surround inhibition measurements, static circular stimuli of increasing diameters were presented at a contrast of 100. The surround inhibition index was calculated using the formula 1-(R_full_/R_pref_) where R_full_ is defined as the response generated to a spot diameter of 800 micron, and R_pref_ represents the response at optimum spot size^55,56^. To assess the direction selectivity of RGCs, a 200-micron spot moving at 1000 micron/s in eight equally spaced directions was presented. Neurons were considered direction-selective if the vector sum of their spike responses was greater than 0.1^89^.

### Contrast sensitivity

The light-evoked spike activity was measured across different contrasts and included in further analysis only when robust responses were observed across 2–3 stimulus trials. The responses were fit to the Naka–Rushton equation R(C) = R_max_(C^n^/(C^n^+C_50_^n^), where R indicates the response at a given contrast C, R_max_ indicates the maximum response, C_50_ indicates the contrast which evokes a half-maximal response and n is a coefficient proportional to the slope of the curve at C_50_. The C_50_ was used as an estimate of contrast sensitivity. Visual threshold was estimated as a second measure of sensitivity under scotopic conditions. It was calculated as the minimum contrast required to elicit a single spike in at least two trials. For the behaviour experiments, contrast sensitivity was estimated from the C_50_ of the normalized contrast response function, where response was measured as described in the optomotor response section.

### Intravitreal Injections

Mice were placed in an induction chamber and anesthetized with 2.0%–2.5% isoflurane delivered in oxygen. Once anaesthetized, the animal was transferred to a standard nose cone to maintain a stable plane of surgical anaesthesia at 1.5% isoflurane. Surgical depth was continuously monitored and verified by loss of the paw withdrawal reflex before the procedure began. The periocular area of both eyes was carefully sanitized with cotton swabs saturated with a topical antiseptic solution (Hypoclin). To prevent corneal desiccation during the procedure, a lubricating ophthalmic solution containing 0.4% w/v polyethylene glycol 400 and 0.3% w/v propylene glycol (Requivisc) was applied topically. Intravitreal injections were performed using a glass micropipette, and the injection pressure was modulated with tubing connected to a 1ml syringe. The micropipette tip was inserted at the limbus, at approximately a 45-degree angle relative to the corneal surface, avoiding damage to the lens. A total volume of 1.5–2.0 µL of either PBS or strychnine 100 μM + Gabazine 1 mM was slowly injected into the vitreous chamber of each eye. Following delivery, the micropipette tip was held in place within the vitreous cavity for 60 seconds to ensure fluid equilibration and prevent post-withdrawal reflux. Throughout the procedure, the animal was placed on a heating pad to maintain its body temperature, and it was returned to its home cage following recovery.

### Immunohistochemistry

Animals were euthanized with an overdose of isoflurane followed by cervical dislocation. The induced diabetic animals were euthanized 13 weeks after STZ injection, the Akita animals were euthanized at 17 weeks of age, and the intravitreally injected animals were euthanized 3 – 7 days post-injection. Eyes were removed using curved forceps and immersed in 4% paraformaldehyde (in PBS) for 10 minutes. Under a dissecting microscope, the eyeballs were hemisected in PBS, and the posterior eyecups were again fixed in 4% paraformaldehyde for 60 minutes at 4°C. The eye cups were immersed in 30% sucrose at 4°C overnight, embedded in the recommended medium, and sectioned at 20 µm thickness using a cryostat (Leica CM1950). These sections were mounted onto glass slides pre-coated with 2% (w/v) APES and stored at-20°C.

Retinal sections were rinsed with PBS and incubated in a blocking buffer (1% BSA, 3% normal goat or donkey serum, and 0.3% Triton X-100 in PBS) for 1 hour at room temperature, followed by overnight incubation with the primary antibodies (Table 1) at 4 °C in a humidified chamber. After washing with PBS, sections were incubated with the secondary antibody for 1 hour at room temperature in the dark. The sections were then washed again with PBS, counterstained with DAPI (1 µg/ml), and mounted in Vectashield medium before being covered with glass coverslips. Imaging was performed using a confocal microscope (Olympus Fluoview 3000) or STED microscope (STEDYCON, Abberior Instruments). The acquired images were adjusted for display purposes using ImageJ. For most analyses, images were acquired from the central retina, defined as the region within ∼ 1,000 µm of the section midpoint. For some analyses, mid-peripheral regions (400 µm from the outer edge to ∼1,000 µm from the center) were also imaged. Two retinal sections were imaged from each animal.

**Table 1:** List of antibodies.

| **Antibody/probe** | **Manufacturer** | **Catalog No.** | **Host** | **Dilution** |
|---|---|---|---|---|
| GFAP | Abclonal | A14673 | Rabbit | 1 in 500 |
| PKCα | Abclonal | A13342 | Rabbit | 1 in 500 |
| PKCα | Santa Cruz | SC8393 | Mouse | 1 in 500 |
| Secretagogin | Biovendor | RD181120100 | Rabbit | 1 in 200 |
| M-Opsin | Sigma-Aldrich | AB5405 | Rabbit | 1 in 500 |
| Rhodopsin | Sigma-Aldrich | MAB5356 | Mouse | 1 in 500 |
| PSD-95 | Enzo | ADI-VAM-PS001-E | Mouse | 1 in 800 |
| Peanut agglutinin-Fluorescein | Vector Labs | FL-1071-5 | NA | 1 in 500 |
| Syntaxin-1 | Sigma-Aldrich | S1172-05ML | Rabbit | 1 in 1000 |
| MnSOD | Cloud Clone | PAB083Mu01 | Rabbit | 1 in 500 |
| cFos | Abclonal | A24620 | Rabbit | 1 in 1000 |
| Anti-rabbit Alexa Fluor 488 | CST | 4412 | Goat | 1 in 500 |
| Anti-mouse Alexa Fluor 488 | CST | 4408S | Goat | 1 in 500 |
| Anti-rabbit Alexa Fluor 594 | CST | 8889 | Goat | 1 in 500 |
| Anti-goat Alexa Fluor 594 | Thermo Fisher | A11058 | Donkey | 1 in 500 |
| Anti-rabbit Alexa Fluor 488 | Thermo Fisher | A21206 | Donkey | 1 in 500 |
| Anti-mouse STAR ORANGE | Abberior | STORANGE-1001 | Goat | 1 in 200 |
| Anti-rabbit STAR RED | Abberior | STRED-1002 | Goat | 1 in 200 |

### TUNEL Assay

Sections were rinsed with PBS and permeabilized using 0.2% Triton X-100 prepared in PBS. For positive controls, sections were pre-incubated with 300 units of DNase (Promega). TUNEL staining was then carried out following the manufacturer’s protocol for frozen sections (Promega DeadEnd™ Fluorometric TUNEL System). Finally, the sections were counterstained with DAPI (1 µg/ml) and mounted before imaging.

### Image analysis

The RBC terminals were isolated using an intensity threshold from 100X magnified images from the confocal/STED microscope, and the size of the particles was measured using the Analyze Particles function. Only particles between 1-40 µm² were considered for further analysis, as manual inspection indicated that individual terminals were in this range. Two sections per animal were analysed. For the MnSOD experiments, the fluorescence intensity within individual terminals was quantified by first processing RBC images using the Cellpose^90^ tool to generate segmentation masks, which were converted into ROIs and exported. These ROIs were then loaded into ImageJ and overlaid onto the corresponding MnSOD images to measure the mean fluorescence intensity within each terminal. MnSOD fluorescence in the RBC terminal, relative to the surrounding IPL region, was used for further analysis.

The length of M-opsin-labelled cone outer segments was estimated from retinal cross-sections imaged at 40X magnification. After isolating the outer segments using an intensity threshold in ImageJ, the outer segment length was estimated using CT-FIRE^91^. The length of the rhodopsin-expressing segments was estimated by a polygon enclosing the rhodopsin-stained region divided by the length of the region evaluated. All final values were calculated by averaging data from two images acquired from two different retinal sections per animal.

For cFos fluorescence quantification, retinal cross sections imaged at 40× magnification were analyzed. In these experiments, analysis was limited to the INL region abutting the ONL, which largely houses the RBCs and CBCs. cFos intensity was estimated in each nucleus after segmenting the DAPI staining using Cellpose. Fluorescence was background-normalized using the cFos fluorescence in the IPL. The position of the soma/nuclei along the INL was estimated using the Y-centroids of DAPI and PKCα ROIs. The positions were normalized to the median of the distribution of the DAPI to ensure alignment of the distribution across samples. Nuclei at positions less than 5 times the median accounted for 80% of the RBCs and were considered RBCs. While nuclei at positions greater than 5 times the median were considered CBCs. Mean cFos fluorescence was then calculated for RBCs and CBCs in control and diabetic animals.

### Statistical Testing

All population data are presented as mean ± SEM and are accompanied by the number of samples in the figure legend. Student’s t-tests were performed to compare different groups and were unpaired unless otherwise specified. Group differences were considered statistically significant when p ≤ 0.05.

## Acknowledgements

We thank Prof. Malcolm Slaughter for commenting on the manuscript. All the imaging experiments were acquired using the confocal microscope housed in the Advanced Bioimaging Facility funded by DST-FIST (SR/FST/LS-II/2020/552(C)). We thank DSS Imagetech for graciously allowing us to use the STED microscope (STEDYCON, Abberior Instruments) and for providing the secondary antibodies needed for STED microscopy.

## Funding statement

This work was supported by a DBT/Wellcome Trust India Alliance Fellowship (IA/I/21/2/505933), a DBT biomedical sciences research grant (BT/PR59281/BMS/85/1462/2025), an ANRF startup research grant (SRG/2024/002505) and an IIT Madras New Faculty Seed Grant (IP/22-23/0847/BT/NFSC/008986) awarded to SS, a CSIR-HRDG fellowship awarded to SAP, and UGC Research Fellowships awarded to PP and LKP.

## Author Contributions

Conceptualization: SS, SAP, NPS and PP. Data Curation, Investigation, and Methodology: SAP, NPS, PP, and HK. Formal analysis and validation: SS, SAP, NPS, PP, HK, AS, LKP and SK. Funding acquisition and Supervision: SS. Visualization: SS, SAP, NPS, PP, HK, AS, and LKP. Writing – original draft: SS and SAP. Writing – review and editing: SS, SAP, NPS, PP, HK, AS, and LKP.

## Conflict of Interest disclosure

The authors report no conflict of interest

## Data availability statement

The data presented in this study will be provided upon request by the corresponding author.

## Supplementary Figures

**Figure S1:**
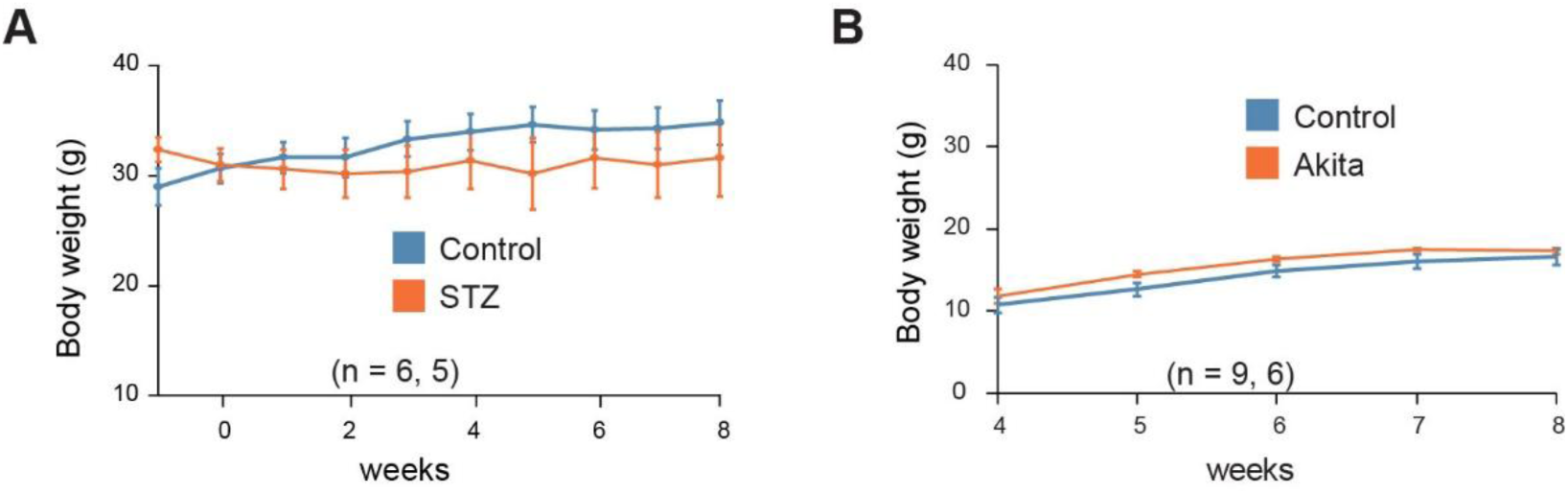
Monitoring body weight in diabetic animal models. (A) Line graph depicting body weight in control and STZ-induced diabetic mice over the time course of the experiment. Body weight was monitored in a subset of the animals shown in Figure 1C, which developed diabetes. Body weight did not change significantly in the diabetic group over the course of the experiment (n = 6 control and 5 diabetic mice). (B) Same as in panel A, but for Akita diabetic animals shown in Figure 1D. Similar to the STZ-injected animals, the diabetic group showed no significant change in body weight over the monitored time period. (n = 9 control and 6 diabetic mice).

**Figure S2:**
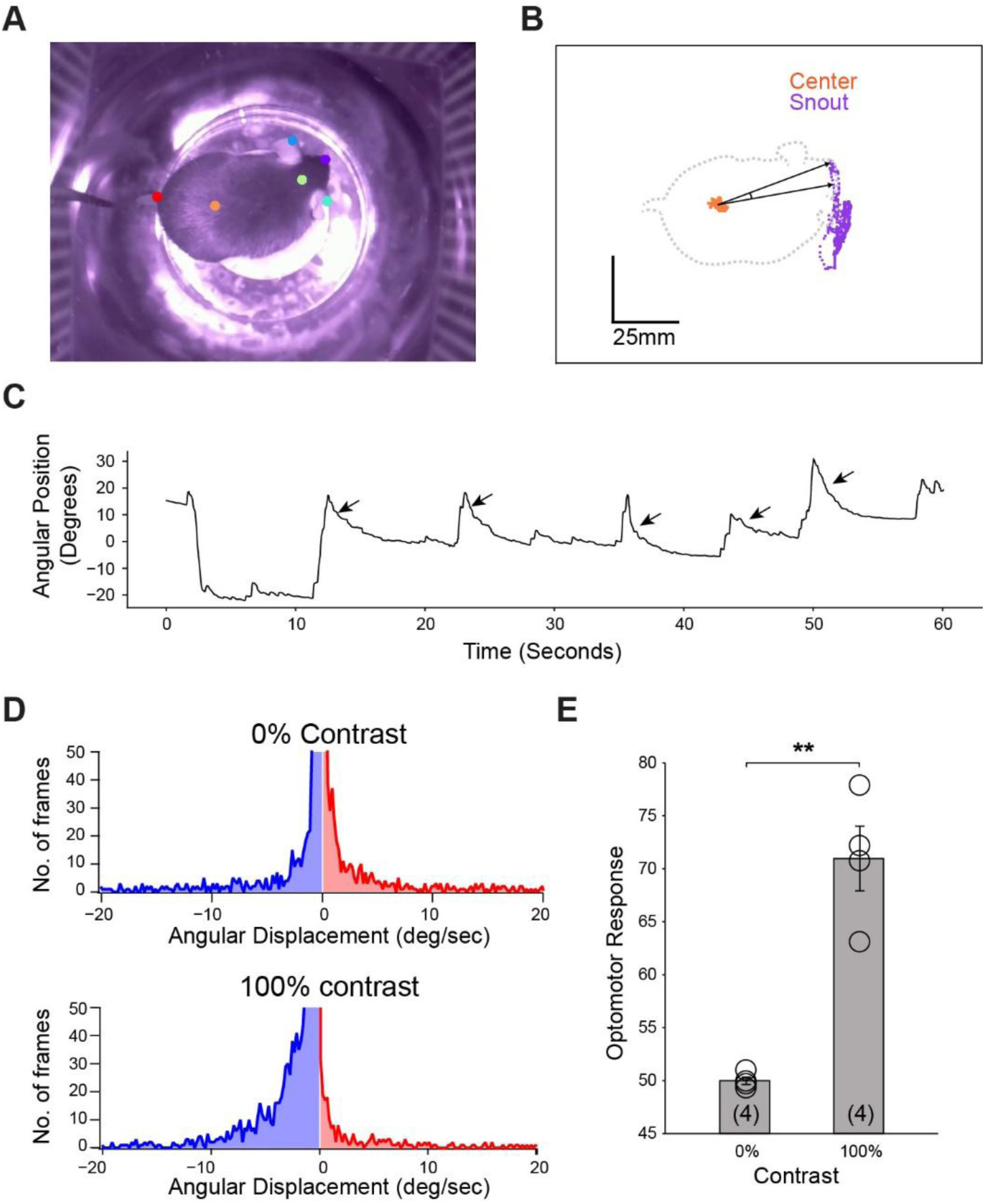
Evaluation of optomotor reflex behaviour in mice. **(A)** A frame from the video capturing the mice performing OMR behaviour. Briefly, the animal is placed on a pedestal in an area, and global motion is simulated using 4 monitors displaying moving black and white gratings with varying contrasts. The videos are processed using DeepLabCut to estimate the position of distinct body parts in each frame. In this frame, the positions of snout (purple), head (green), left ear (blue), right ear (light blue), center (orange) and tail base (Red) are highlighted. **(B)** Spatial coordinates (*x*, *y*) of the snout (purple) and body center (red) estimated across all frames of the recorded behavioral session of 60 seconds at 30fps. From these coordinates, the angular displacement of the snout relative to the center is calculated for each frame. **(C)** Temporal evolution of the snout’s angular displacement in a single video. Small black arrows denote specific head-tracking OMR behavioral events, characterized by slow movements in the direction of stimulus motion followed by a fast reset. **(D)** Frequency distribution of angular displacements of gratings moving in the anticlockwise direction at 0% contrast (uniform grey screen; top) and 100% contrast (bottom). **(E)** A comparison of the Optomotor Response expressed as the percentage of correct head movements (i.e. in the direction of stimulus motion) for 0 and 100% contrast (n = 4, mean ± SEM; **p < 0.01). These responses were normalized for the analysis shown in Figures 1 and 6.

**Figure S3:**
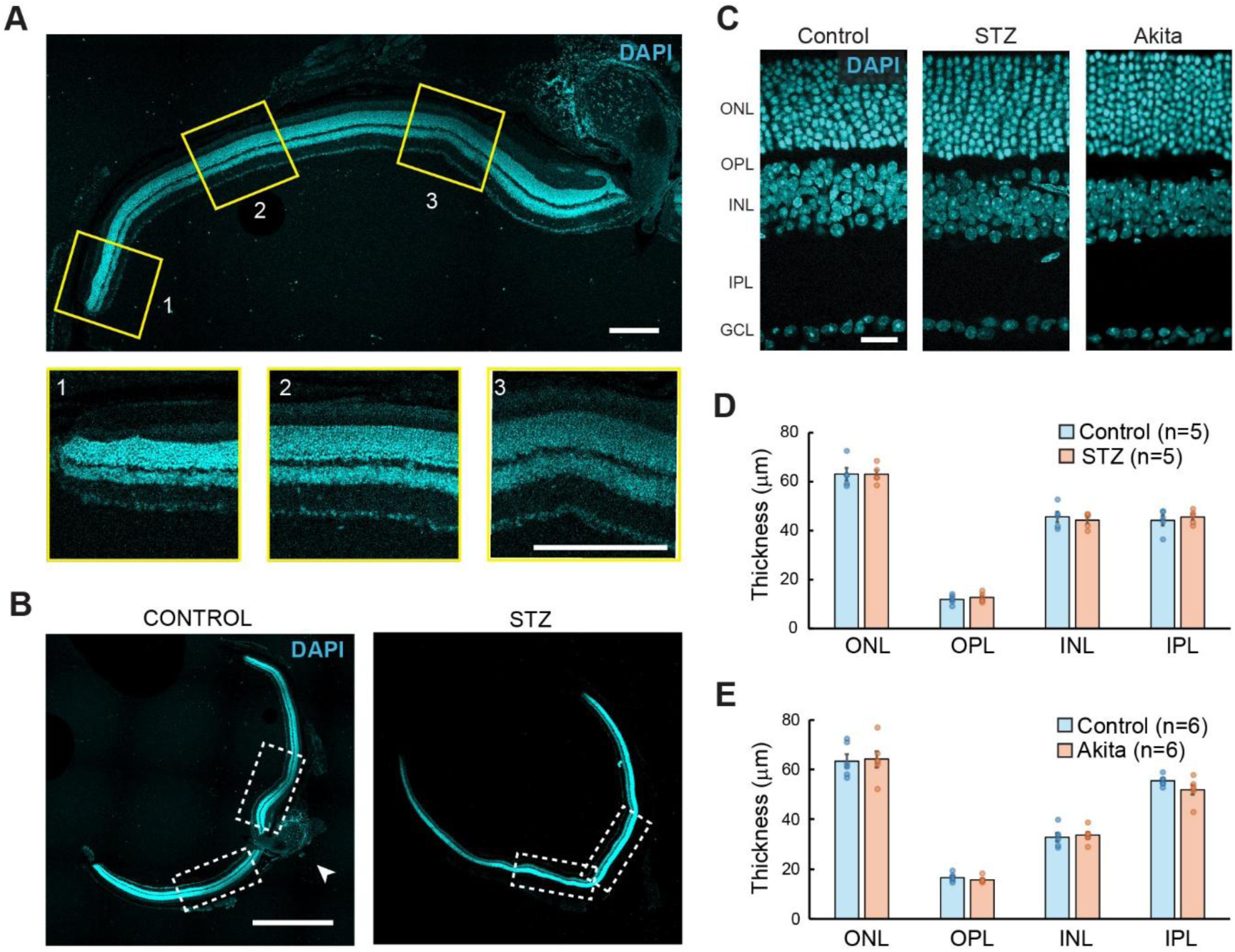
Quantification of retinal layer thickness in control and diabetic mice. **(A)** Stitched 10× images of a control retinal section showing a gradual increase in retinal layer thickness toward the central region. Magnified views of the three yellow insets are displayed below. Scale bar: 200 μm. **(B)** Stitched 10× images of control (left) and STZ-induced diabetic (right) retinal sections. Thickness measurements were performed in areas near the center, as marked by the white boxes. The arrowhead indicates the optic disc, which was excluded from the analysis as it is dominated by ganglion cell axons. Scale bar: 1000 μm **(C)** Retinal sections from control, STZ-injected and Akita mice at 13 weeks post-STZ injection, with nuclear layers labelled in blue. Scale bar: 20 μm. **(D)** Bar graph illustrating the thickness of different layers in control versus STZ-injected mice. There was no significant difference in the thickness of any of the retinal layers between the two groups. Data represented as mean ± SEM, with circles indicating the individual data points (n = 5 mice per group). **(E)** Same as D but comparing retinal thickness between control and Akita retina. There was no significant difference in the thickness of any of the retinal layers between the two groups.

**Figure S4:**
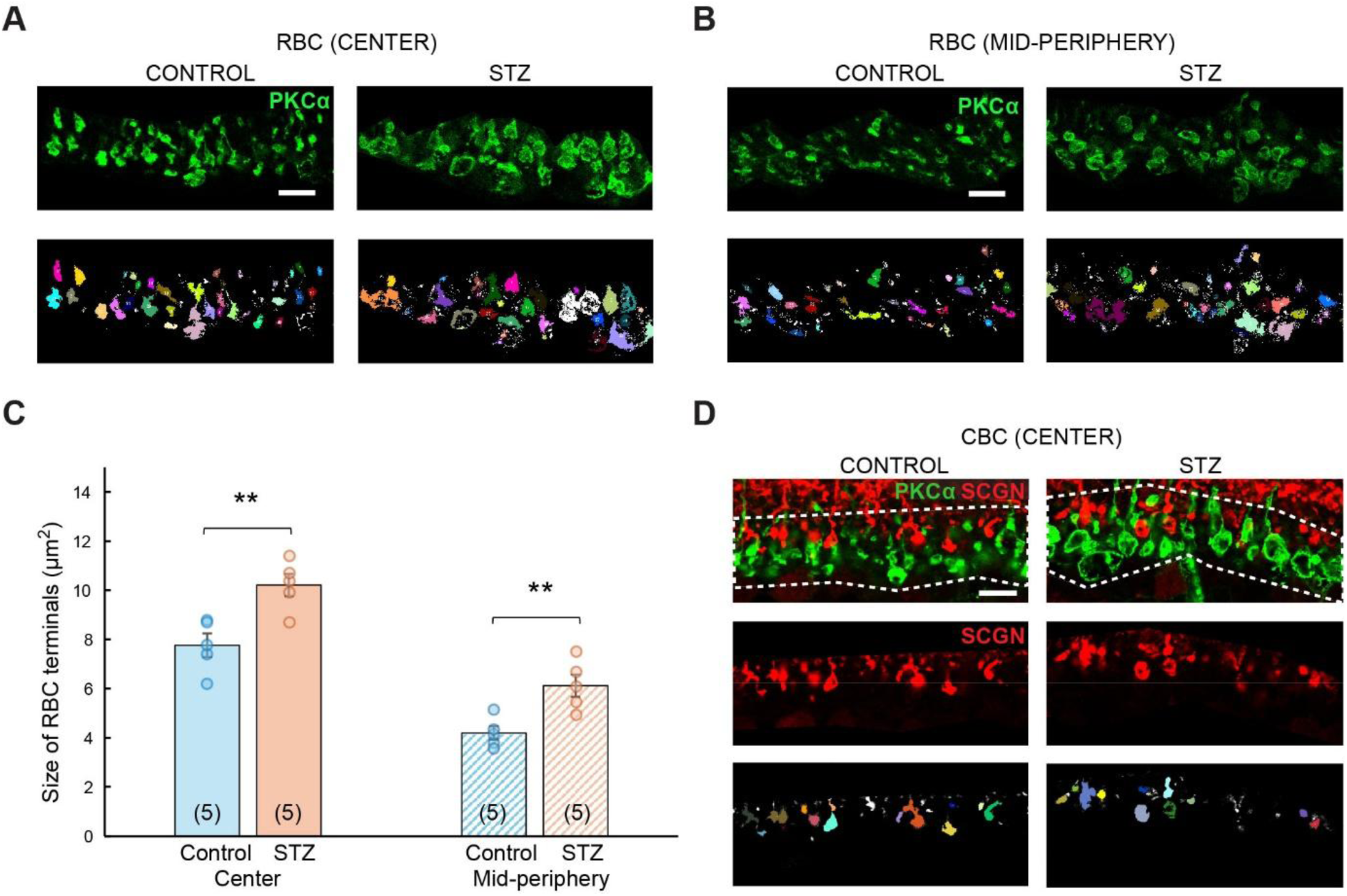
Quantification of axonal terminals of bipolar cells. **(A)** Representative images showing RBC axon terminals (in green) from the central region of control (left) and STZ-injected diabetic (right) retinas. Scale bar: 10 µm. Bottom panels display the identified terminals as ROIs from ImageJ’s Analyze Particles function. **(B)** Same as (A) but showing terminals from the mid-peripheral region. RBC terminals in the mid-peripheral region were smaller than the terminals in the center. **(C)** Bar graph comparing the average size of RBC terminals between control and STZ-injected diabetic groups in both central and mid-peripheral regions. Data represented as mean ± SEM, with individual data points shown as circles (n = 5 mice per group; **p < 0.01). **(D)** Top panel shows CBC terminals (in red) located closest to RBC terminals (in green), likely within strata 4 and 5 of the IPL (outlined by white dashed line) in a control (left) and diabetic (right) retinal section. The middle panel presents the cropped image used for analysis, and the bottom panel displays the segmented output from ImageJ. Scale bar: 10 µm

**Figure S5:**
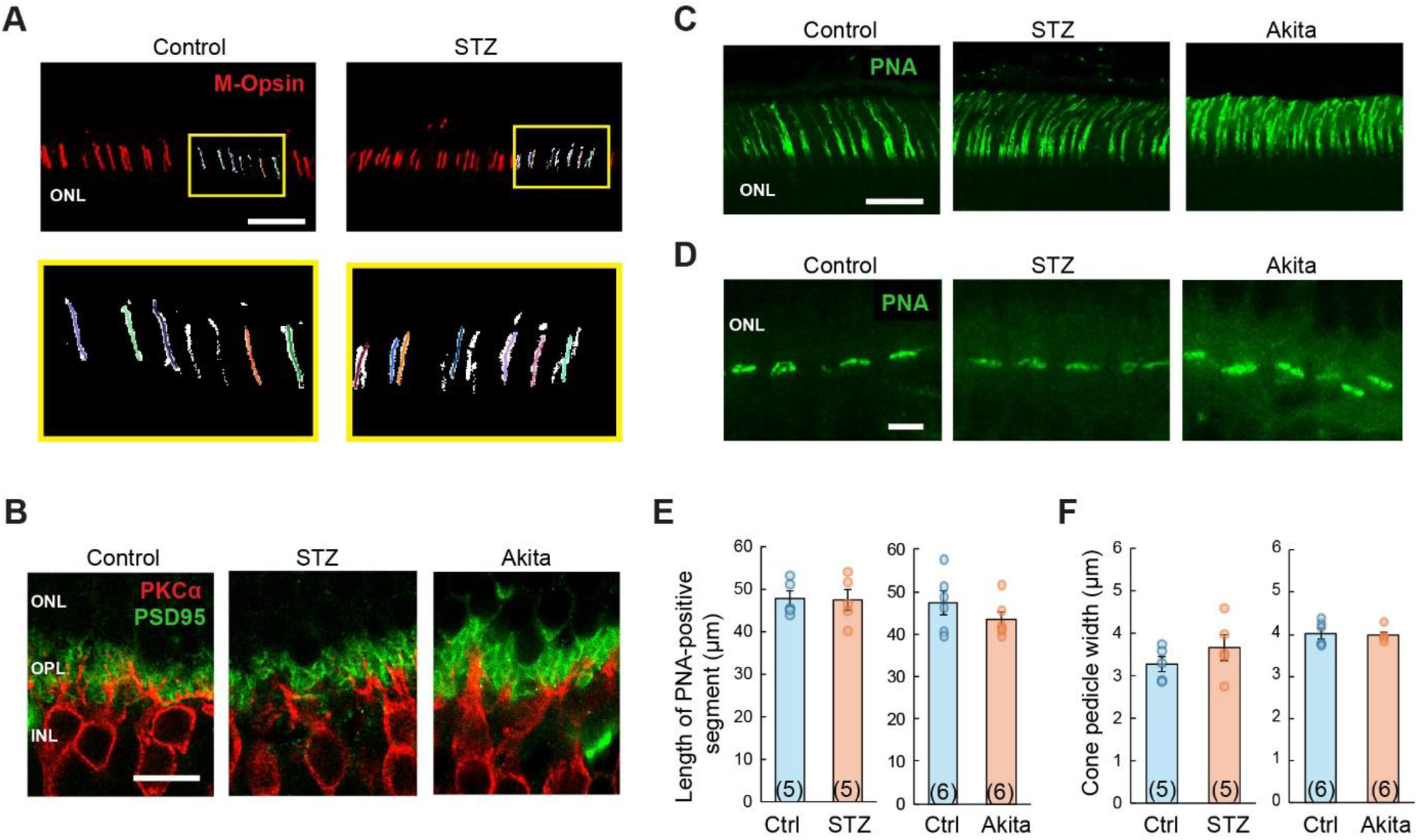
Assessment of rod and cone photoreceptors in diabetic mice. **(A)** Top panel shows M-opsin expression (red) in the outer segments of photoreceptors in control (left) and STZ-injected diabetic (right) mouse retina. Scale bar: 40 µm. The inset displays the CT-FIRE output, where each line represents an individual fragment segmented as a separate structure. A magnified view of the inset is shown in the bottom panel. **(B)** Retinal section from control, STZ-injected and Akita mice, stained with anti-PSD-95 (green) to visualize photoreceptor synaptic terminals and anti-PKCα (red) to label rod bipolar cells. The characteristic horseshoe-shaped photoreceptor terminals are largely unaltered in the diabetic models. Scale bar: 10 µm **(C)** Representative images showing PNA labelling (green) of the cone outer segments in control, STZ-injected and Akita mouse retinas. Scale bar: 40 µm. **(D)** Representative images showing PNA-labeling (green) of the cone pedicles in control, STZ-injected and Akita mouse retinas. Scale bar: 5 µm. **(E)** Bar graphs comparing the length of the PNA-labeled cone outer segments in control and diabetic retinas. Data represented as mean ± SEM, with circles indicating the individual data points. Similar to the M-opsin staining, the length of the PNA stained outer segments was not significantly different between control and diabetic retina. **(F)** Bar graphs comparing cone pedicle width in control and diabetic retinas. The PNA-labeled structures in cone pedicles were quantified by fitting them into ellipses using the Analyze particles function in ImageJ. The major axis of the ellipse was considered as the length of the pedicle. Data represented as mean ± SEM, with circles indicating the individual data points. The PNA-stained cone pedicle width was not significantly different between the control and diabetic groups.

**Figure S6.**
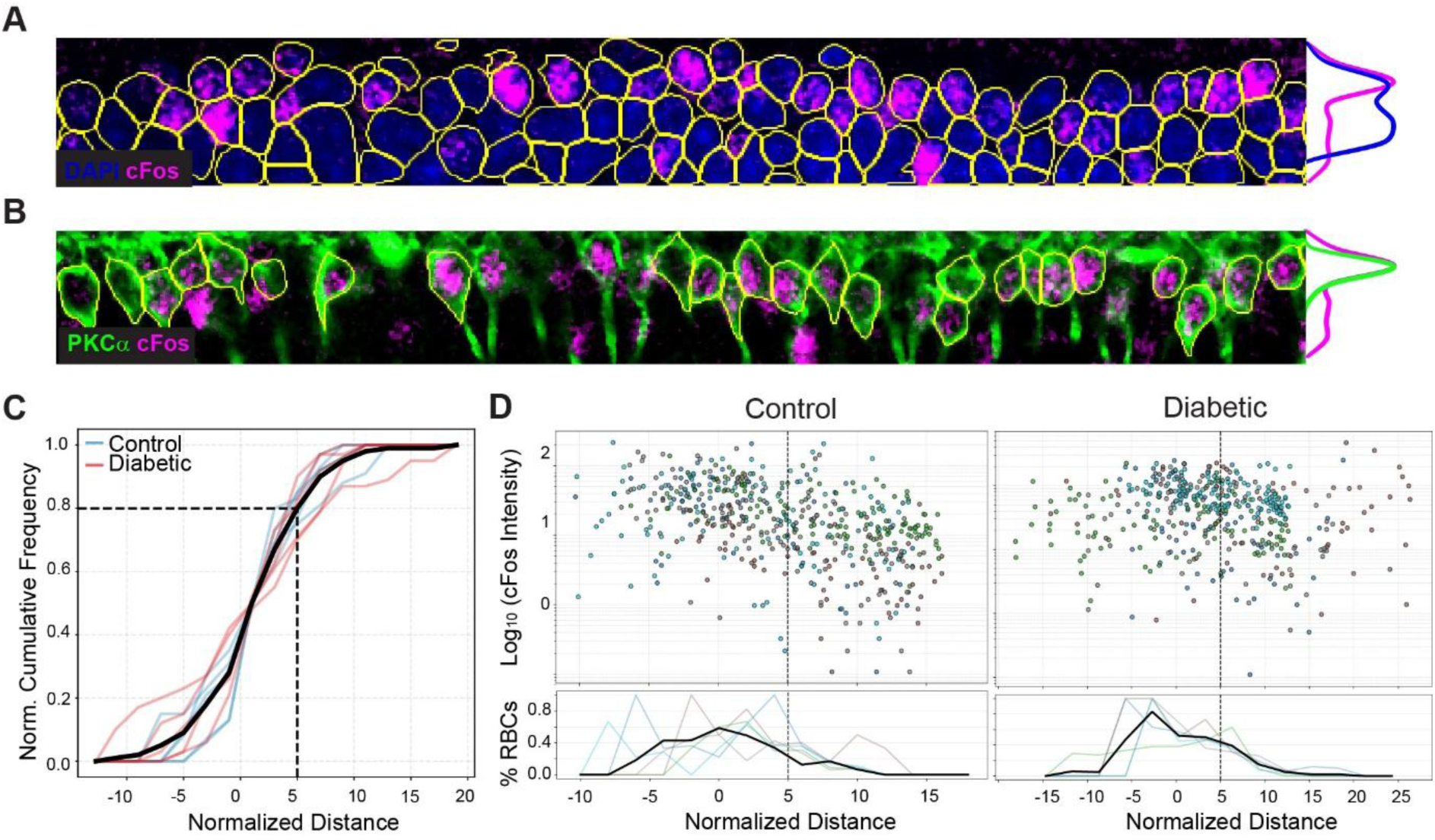
Spatial distribution of c-Fos expression in the bipolar cells along the INL depth. **(A)** Representative retinal image showing DAPI-labelled nuclei (blue) and c-Fos immunoreactivity (magenta). The yellow circles indicate the outlines of nuclei extracted from Cellpose using DAPI staining. The curves on the right indicate the frequency profiles of the distribution of DAPI-positive ROIs and c-Fos fluorescence intensity along the y-axis of the image. **(B)** Same region shown in A, but for PKCα-labelled soma (green) and c-Fos immunoreactivity (magenta). The yellow ROIs indicate the outlines of PKCα-positive somas extracted from Cellpose. The curves on the right depict the frequency distribution of PKCα-positive ROIs and c-Fos fluorescence intensity along the y-axis of the image. **(C)** Line plot showing the normalized cumulative frequency distribution of rod bipolar cells (PKCα-positive ROIs) as a function of normalized distance along the INL. Negative values indicate the soma is closer to the OPL and positive values indicate that the soma is closer to the IPL. Semi-transparent blue and red lines represent individual control and diabetic animals, respectively, while the bold black line represents the mean cumulative frequency distribution across animals. Approximately 80% of rod bipolar cells are located within x = 5. Nuclei at positions lower than x = 5 were considered rod bipolar cells and nuclei higher than this value were assigned cone bipolar cells (Figure 3). **(D)** Scatter plot showing normalized c-Fos fluorescence intensity of each nucleus compared to its position in the INL in control (left) and diabetic (right) retina. The line plots at the bottom show the frequency histogram of rod bipolar cells (PKCα-positive ROIs) relative to the total number of DAPI-positive ROIs. Semi-transparent lines represent individual control and diabetic animals, respectively, while the bold black line represents the mean proportion of rod bipolar cells across animals. The black dashed line at x = 5 indicates the threshold used to distinguish rod bipolar cells and cone bipolar cells. The population averages of these two groups are shown in Figure 3C**, D**.

**Figure S7:**
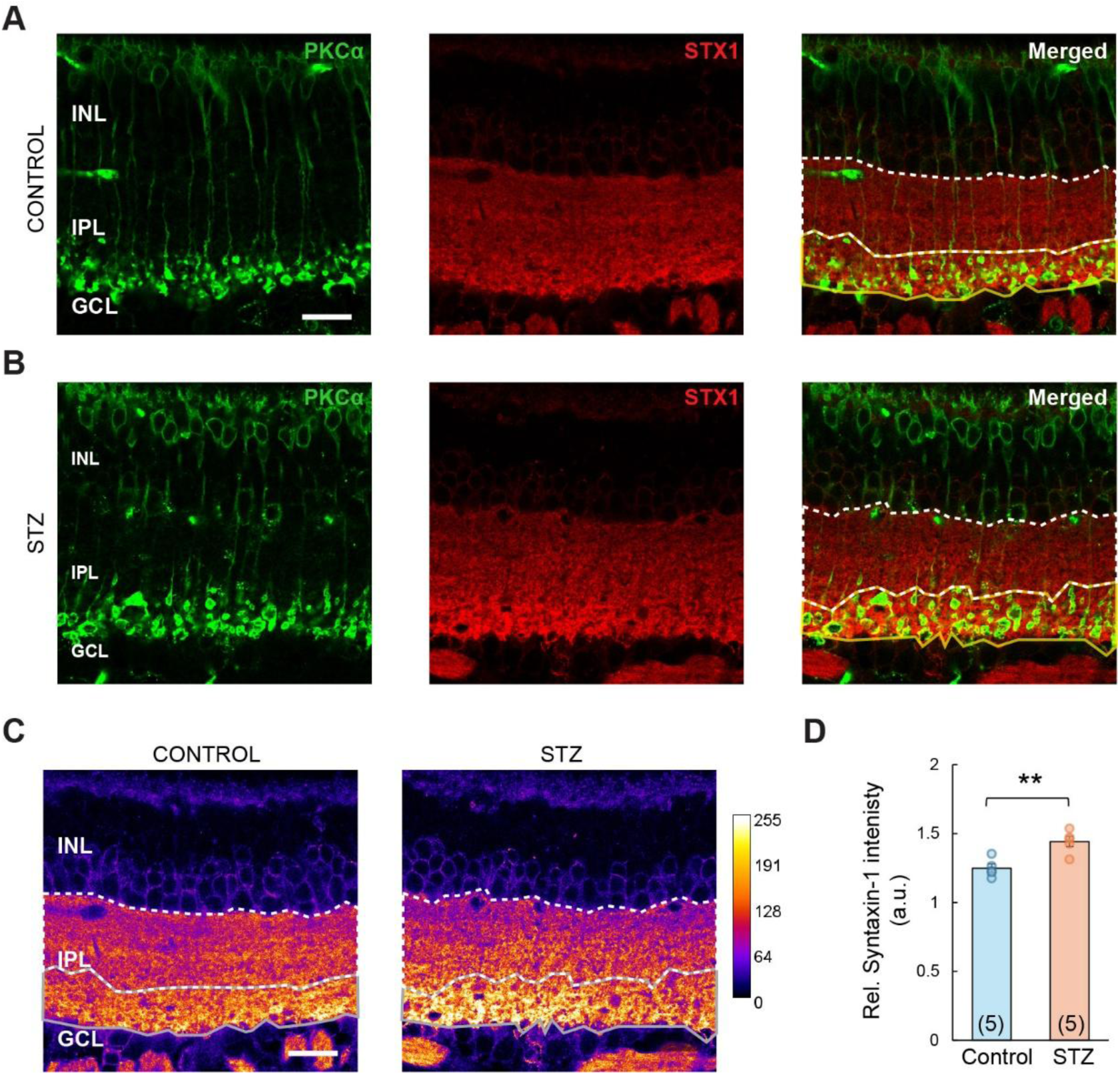
Elevated syntaxin-1 expression in RBC terminal layer suggests altered amacrine cell activity. **(A)** Retinal sections from control mice labelled with anti-PKCα (green) and anti-syntaxin-1 antibody (red). The yellow outline marks the RBC terminal layer, while the white dashed outline denotes the remaining portion of the IPL. Scale bar: 20 µm. **(B)** Same as A, but for STZ-injected diabetic retina **(C)** The Syntaxin-1 labelling shown in A and B, but visualized using the Fire lookup table for enhanced contrast. The accompanying color scale bar indicates grayscale values ranging from 0 to 255, corresponding to increasing fluorescence intensity. **(D)** Bar graph comparing the relative intensity ratios of syntaxin-1 fluorescence between control and STZ-injected diabetic retinas. Data represented as mean ± SEM, with circles indicating the individual data points. The Syntaxin levels in the RBC layer was significantly higher in the diabetic tissue compared to control (n = 5 mice per group; **p = 0.004).

**Figure S8:**
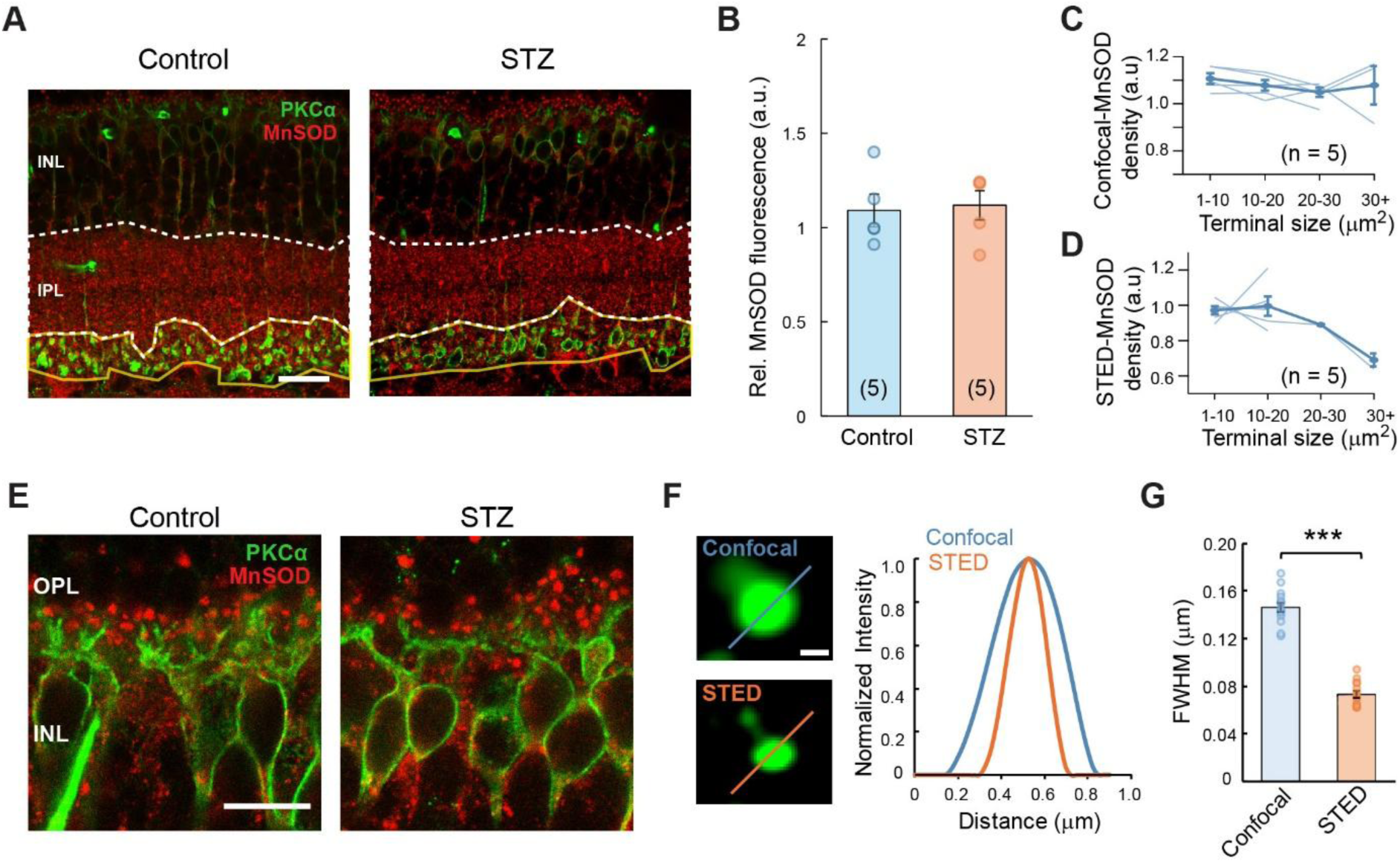
Assessment of mitochondria under hyperglycemic conditions. **(A)** Retinal sections from control (left) and STZ-injected diabetic (right) mice labeled with anti-PKCα (in green) and anti-MnSOD antibody (in red). The yellow outline marks the RBC terminal layer, while the white dashed outline denotes the remaining portion of the IPL. Scale bar: 20 µm. **(B)** Bar graph comparing the relative intensity ratios of MnSOD fluorescence in the RBC layer between control and STZ-injected diabetic retinas. Data represented as mean ± SEM, with circles indicating the individual data points (n = 5 mice per group). **(C)** Line graph showing changes in MnSOD expression across increasing terminal sizes for each control animal (thin lines). The group mean is represented by a solid blue line, with error bars indicating the SEM. Note: Y-axis does not start from zero to better visualize subtle changes. **(D)** Similar to C, but for images collected using STED microscopy **(E)** Retinal section from control and STZ-injected Akita mice, stained with anti-MnSOD (red) and anti-PKCα (green) to visualise MnSOD puncta in OPL. Scale bar: 10 µm **(F)** Representative confocal and STED images of the same puncta (Scale bar: 200 nm). Line intensity profiles were extracted along the indicated line (1 μm) and fitted with a Gaussian function. **(G)** Bar plot shows the Full Width at Half Maximum (FWHM) values obtained from the Gaussian fits shown in F (n = 14 puncta from 2 images). The FWHM of the STED microscope was ∼70nm compared to the confocal ∼140nm (***p<0.001; paired Student’s t-test).

**Figure S9:**
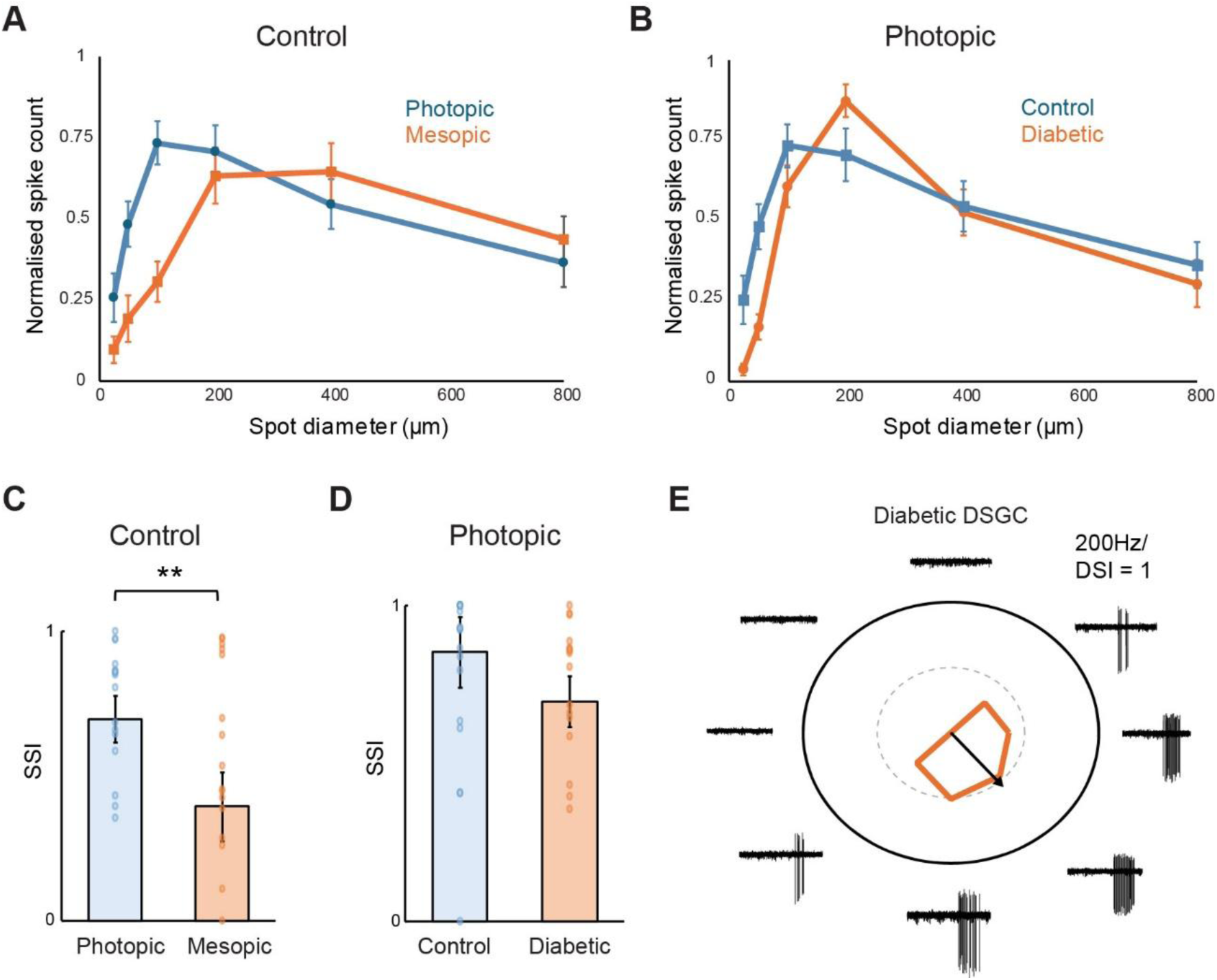
Surround suppression and direction selectivity in diabetic retinal ganglion cells. **(A)** Normalized spike counts of control retinal ganglion cells (n = 16 cells) are plotted against spot diameter under photopic (blue) and mesopic (orange) conditions. Responses were normalized to the maximal response of each cell. Data represented as mean ± SEM. **(B)** Normalized spike counts of retinal ganglion cells (n = 16 cells) are plotted against spot diameter under photopic conditions in control (blue) and diabetic (orange) retinas. Responses were normalized to the maximal response of each cell. Data represented as mean ± SEM. **(C)** Comparison of the surround suppression index (SSI) of retinal ganglion cells in photopic and mesopic conditions in control retina (n = 16 cells). Retinal ganglion cells exhibited significantly stronger surround suppression under photopic than under mesopic conditions (**p = 0.004; paired t-tests). Individual data points are shown as open circles, and bars represent the mean ± SEM. **(D)** Comparison of the surround suppression index (SSI) between control and diabetic retinal ganglion cells under photopic conditions (n = 16 cells). No significant difference was observed in surround suppression between control and diabetic groups (p = 0.708; unpaired t-test). Individual data points are shown as open circles, and bars represent the mean ± SEM. **(E)** Polar plot illustrating the direction selectivity of a representative retinal ganglion cell from a diabetic retina (n = 1). Spike responses elicited by a spot of light moving in eight different directions are shown as representative traces around the polar plot. The vector indicates the preferred direction of the cell, with its length representing the direction selective index.

**Figure S10:**
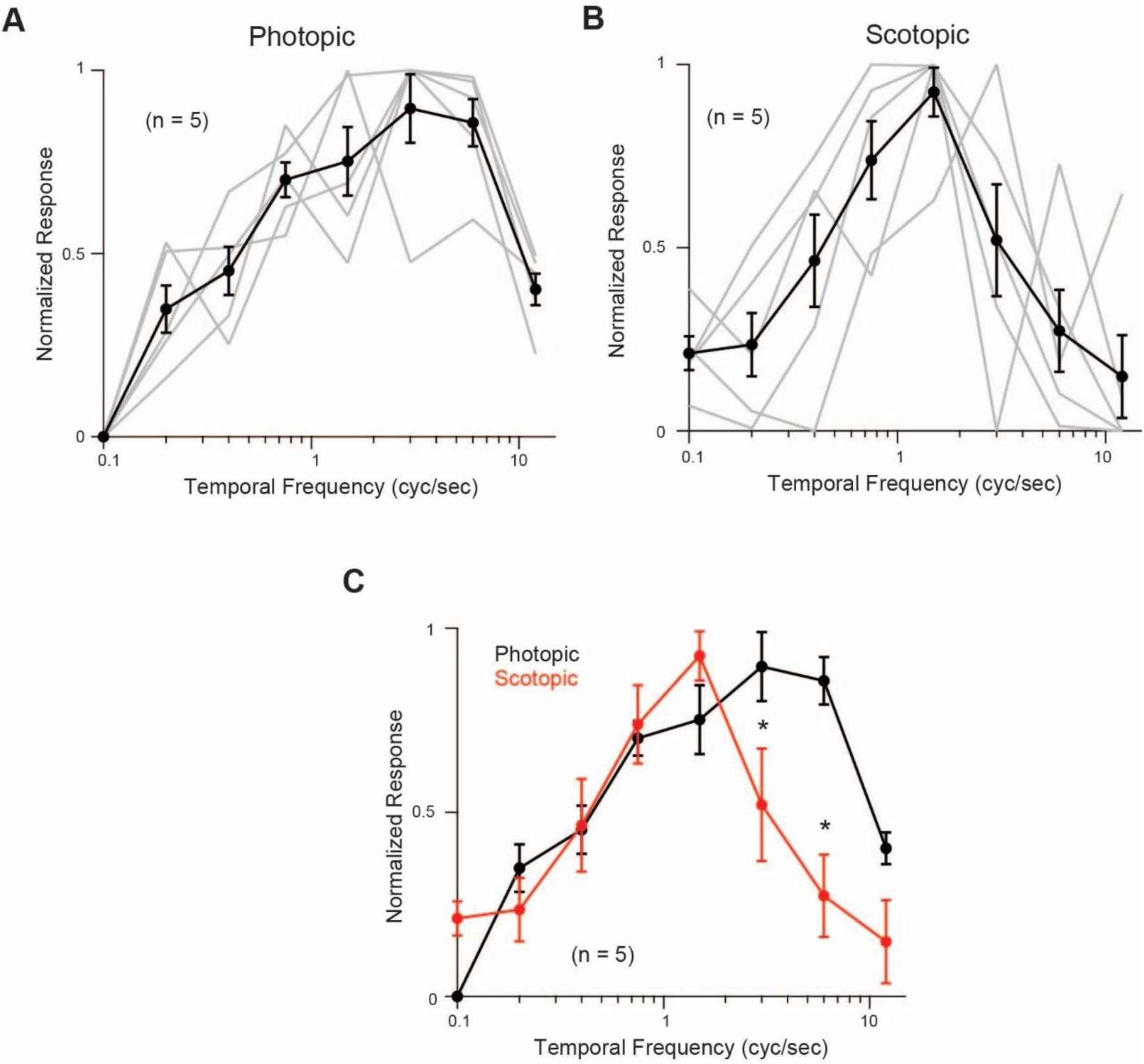
Temporal Frequency Tuning Profiles Under Photopic and Scotopic Conditions. **(A)** Normalized OMR response for sinusoidal grating of a fixed spatial frequency (0.2 cycles/degree) but with varying temporal frequencies (0.1 to 12 Hz) under photopic conditions. The circles and the black line indicate the population response, while the grey lines represent individual responses (n = 5). Data represented as mean ± SEM. **(B)** Similar to A, but under scotopic conditions. **(C)** Overlay of the photopic (black) and scotopic (red) responses shown in A and B, respectively. The response at higher temporal frequencies was significantly higher in photopic conditions compared to scotopic conditions (*p < 0.05; paired t-test).

## Notes

### Competing Interest Statement

The authors have declared no competing interest.

